# Pan-genome Analysis in Sorghum Highlights the Extent of Genomic Variation and Sugarcane Aphid Resistance Genes

**DOI:** 10.1101/2021.01.03.424980

**Authors:** Bo Wang, Yinping Jiao, Kapeel Chougule, Andrew Olson, Jian Huang, Victor Llaca, Kevin Fengler, Xuehong Wei, Liya Wang, Xiaofei Wang, Michael Regulski, Jorg Drenkow, Thomas Gingeras, Chad Hayes, J. Scott Armstrong, Yinghua Huang, Zhanguo Xin, Doreen Ware

## Abstract

*Sorghum bicolor*, one of the most important grass crops around the world, harbors a high degree of genetic diversity. We constructed chromosome-level genome assemblies for two important sorghum inbred lines, Tx2783 and RTx436. The final high-quality reference assemblies consist of 19 and 18 scaffolds, respectively, with contig N50 values of 25.6 and 20.3 Mb. Genes were annotated using evidence-based and *de novo* gene predictors, and RAMPAGE data demonstrate that transcription start sites were effectively captured. Together with other public sorghum genomes, BTx623, RTx430, and Rio, extensive structural variations (SVs) of various sizes were characterized using Tx2783 as a reference. Genome-wide scanning for disease resistance (R) genes revealed high levels of diversity among these five sorghum accessions. To characterize sugarcane aphid (SCA) resistance in Tx2783, we mapped the resistance region on chromosome 6 using a recombinant inbred line (RIL) population and found a SV of 191 kb containing a cluster of R genes in Tx2783. Using Tx2783 as a backbone, along with the SVs, we constructed a pan-genome to support alignment of resequencing data from 62 sorghum accessions, and then identified core and dispensable genes using this population. This study provides the first overview of the extent of genomic structural variations and R genes in the sorghum population, and reveals potential targets for breeding of SCA resistance.

## INTRODUCTION

*Sorghum bicolor*, the fifth most economically important cereal, forage, and bioenergy crop in the world (Ordonio et al., 2016), is known as the “camel of the grass family” due to its high heat and drought tolerance. Hence, accelerating crop improvement in sorghum is key to ensuring global food and energy security in the context of climate change (Morris et al., 2013). Moreover, its small and compact genome relative to other C4 grasses makes it an excellent model for genomic studies (Paterson et al., 2009). Previous population genomic and genome-wide association studies of agroclimatic traits in sorghum provided a basis for crop improvement through marker-assisted breeding and genomic selection (Morris et al., 2013). In addition, whole-genome sequencing of different sorghum lines spanning a wide range of geographic origins, indicated that sorghum offers underdeveloped genetic resources that are unique among the major cereals (Mace et al., 2013).

Previous studies of sorghum diversity were mainly limited to single-nucleotide polymorphisms (SNPs) and small insertions/deletions (indels) called from short-read sequencing technologies (Mace et al., 2013; Brenton et al., 2016; Zarrei et al., 2015; Saxena et al., 2014). Recent development of long-read sequencing technologies have brought genomics studies into a new era for many species, including maize (Jiao et al., 2017), rice (Zhang et al., 2016), *Arabidopsis* (Michael et al., 2018), and sorghum (Deschamps et al., 2018). However, the diversity within a species cannot be represented by a single reference genome, and this realization has led to the spread of the pan-genome concept. A pan-genome represents the genomic diversity of a species and includes core genes found in all individuals, as well as variable genes that are absent from some individuals (Bayer et al., 2020). In addition, genomic structural variations (SVs), including deletions, insertions, inversions, and duplications, represent an important source of genetic diversity (Alkan et al., 2011; Baker, 2012; Zhang et al., 2015), and such variations have been implicated in several human diseases (Sebat et al., 2007; Stefansson et al., 2008). In plants, SVs are associated with variations in phenotypes including leaf size (Horiguchi et al., 2009), fruit shape (Xiao et al, 2008), and aluminum tolerance (Maron et al., 2013). Accurate identification and characterization of large SVs will require long reads and high-quality genome assemblies. Long reads are advantageous for SV calling because they can span repetitive or those that are problematic for other reasons (Mahmoud et al., 2019). One class of gene-space structural variants are the disease-related nucleotide-binding site plus leucine-rich repeat (NBS-LRR) genes, which in sorghum have higher diversity than non–NBS-LRR genes (Mace et al., 2014). Several NBS-LRR genes have also been identified in *Arabidopsis* (Meyers et al., 2003) and rice (Stein et al., 2018; Zhou et al., 2004).

Here, we report the development of two new high-quality chromosome-level reference assemblies of sorghum: Tx2783, a widely utilized pollinator parent with sugarcane aphid (SCA) resistance, and RTx436, another widely utilized pollinator parent with known general combinability. We used the new reference genomes and three publicly available accessions, BTx623 (McCormick et al., 2018), RTx430 (Deschamps et al., 2018), and Rio (Cooper et al., 2019), to characterize SVs. Using a recombinant inbred line (RIL) population between Tx2783 and BTx623, we identified an SV on chromosome 6 that segregates with a QTL identified for SCA tolerance. In addition, we constructed a sorghum pan-genome using TX2783 as the backbone, along with insertions called from the four complete genomes, and used it to identify core and dispensable genes based on resequencing data from 62 sorghum accessions. Our results provide new insights into sorghum population diversity, as well as potential targets for breeding sorghum for high sugarcane aphid tolerance.

## RESULTS

### Chromosome-level Genome Assembly of Two Sorghum Inbred Lines

To characterize large SVs in the sorghum population and provide support for modern breeding, we selected two important sorghum inbred lines, Tx2783 and RTx436, for genome construction. Line Tx2783 (PI 656001) was originally bred for resistance to sorghum greenbug (*Schizaphis graminum*) and also exhibits high SCA resistance (Tetreault et al., 2019). The other line, RTx436 (PI 561071), is a widely adapted pollinator parent used in the development of high-yielding hybrids. RTx436 is also commonly used in the development of traditional food-grade sorghum hybrids with white grain color and tan glumes, which are suitable traits for many food processors. These two sorghum lines, Tx2783 and RTx436, were sequenced using PacBio CLR technology to coverage of 76× (reads N50=23.1 kb) and 61× (reads N50=24.7 kb), respectively. The assembly effort generated contigs with N50 lengths of 25.6 Mb for Tx2783 and 20.3 Mb for RTx436. In addition, we generated Bionano molecules for Tx2783 and RTx436 that yielded genome maps of 721.504 Mb and 723.680 Mb, respectively, with N50 lengths of 36.987 Mb and 37.781 Mb (Supplemental Table S1). The chromosomes of these two genomes were constructed using hybrid scaffolds generated from Bionano genome maps. Most of the chromosomes consisted of two scaffolds (Fig. 1A, Table 1). These two genomes were of higher quality than the published sorghum reference genomes of BTx623 (McCormick et al., 2018) and RTx430 (Deschamps et al., 2018), with 15-fold and 7-fold higher average sequence continuity, respectively, and a total assembly size that is much closer to the estimated size of the sorghum genome, ~730 Mb. Gene space assessment using BUSCO confirmed the completeness of the Tx2783 and RTx436 genome assemblies (Fig.1B).

**Figure 1.**
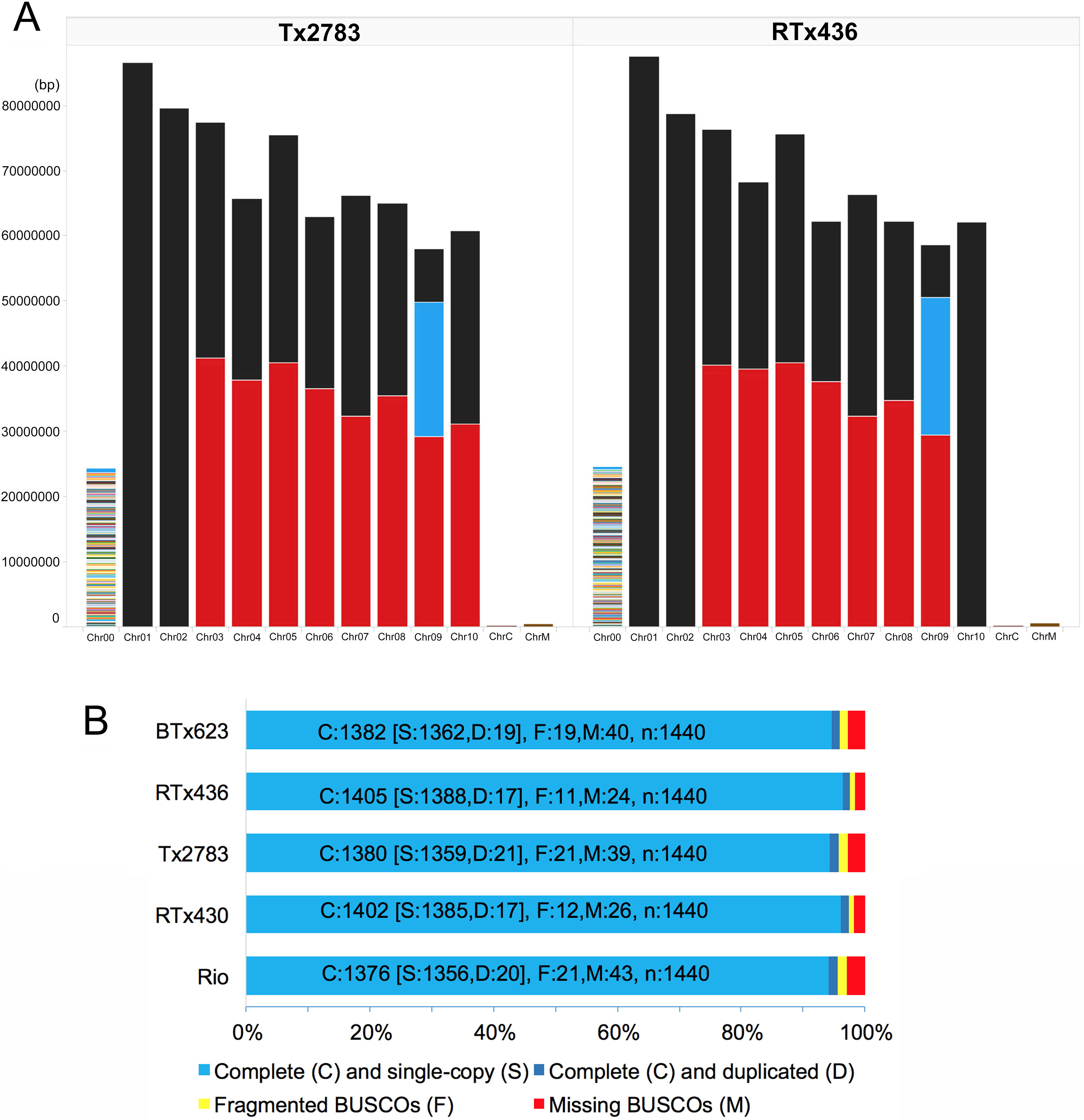
Number of scaffolds per chromosome and BUSCO assessment. **(A)**. Number of scaffolds per chromosome for the Tx2783 and RTx436 genome assemblies. Each colored bar represents a scaffold. Most scaffold breaks are occurred at the centromeres. Chr00: Contigs unplaced into the chromosome. ChrC: Chlotoplast sequence. ChrM: Mitochondria sequence. **(B)**. BUSCO assessment of the indicated sorghum genomes.

### Transposable Element and Gene Annotation of the Tx2783 and RTx436 Genomes

Transposon elements (TEs) were annotated using EDTA (Ou et al., 2019), with 68.97%, 69.34%, and 69.35% of genome sequences annotated as TEs in Tx2783, RTx436, and BTx623, respectively. The majority (53.23%, 53.29%, and 53.19%) of TEs in each genome were annotated as retrotransposons. Of those, LTR-Gypsy was the most abundant, representing 36.63%, 37.20%, and 37.98% of TEs in the Tx2783, RTx436, and BTx623 genomes, respectively (Supplemental Table S2). In addition, we also identified 10984 (79.5%), 11207 (79.7%), and 10746 (79.2%) intact LTRs in the Tx2783, RTx436, and BTx623 genomes, respectively.

Gene calling was performed using a hybrid approach with evidence-based and *de novo* gene predictors (see Methods), and then filtered based on annotation evidence distance (AED) score (Campbell et al., 2014) and homology to a maize, *Brachypodium*, rice, or *Arabidopsis* protein sequences obtained from Gramene release 62 (Tello-Ruiz et al., 2020). Ultimately, this approach generated a total of 29,612 and 29,265 protein-coding genes and 4,205 and 3,478 non-coding genes in the Tx2783 and RTx436 genomes, respectively (Supplemental Table S3). However, the RTx436 annotation predicted slightly longer genes with a higher intron count; accordingly, Tx2783 had slightly more single-exon genes than RTx436. In addition, CDSs, peptides, and 5’ and 3’ UTRs were longer in Tx2783 than in RTx436. To assess the quality of the 5’ transcriptional start sites, root and shoot tissues from Tx2783 and RTx436 were collected for RAMPAGE (RNA annotation and mapping of promoters for analysis of gene expression) assays (Batut et al., 2013). This analysis identified 228,249 and 300,633 high-confidence peaks from Tx2783 and RTx436 respectively. The narrow distribution of RAMPAGE signals (Fig. 2A and 2B) over annotated loci confirmed our very high specificity for true TSSs in both root and shoot tissues from the Tx2783 and RTx436 genomes. The high-confidence peaks overlapped in 27,135 and 26,432 genes in Tx2783 and RTx436 respectively, indicating that the two genomes were well annotated.

**Figure 2.**
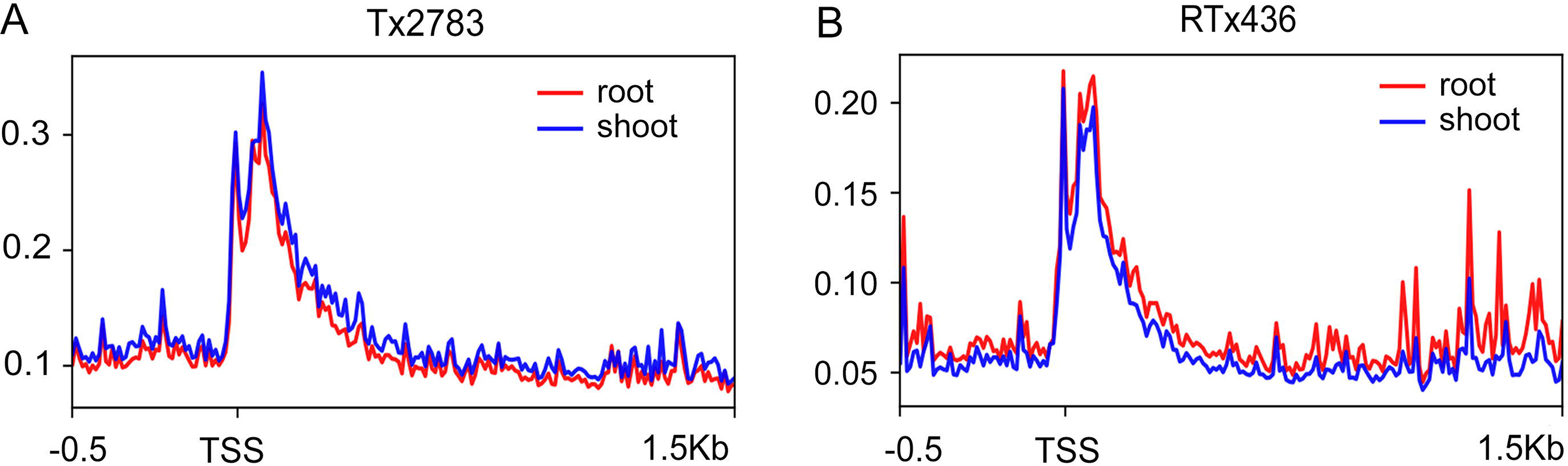
RAMPAGE signal enrichment around TSSs in Tx2783 (A) and RTx436 (B) in root and shoot tissues.

### Characterization of Genetic Variations in Five Sorghum Reference Assemblies

We initially assessed large-scale structural variations between the accessions by comparing alignments between the BTx623 Bionano map and the sorghum v3.1 reference sequence. This comparison revealed several large inversions (Fig. 3A), which were also found in the alignment between Tx2783, RTx436, and the v3.1 reference (Fig. 3B and 3C) but were absent in comparisons between Tx2783 or RTx436 and the BTx623 Bionano map (Fig. 3D and 3E) and between Tx2783 and RTx436 (Fig. 3F), suggesting potential artifacts in the public BTx623 reference assembly. A list of these SVs from different alignments is provided in Supplemental Data S1. A total of 36 large inversions (>100kb) were detected in the two lines in the same area (Supplemental Table S4). The pseudomolecules of the reference genome BTx623 was constructed by genetic maps (McCormick et al., 2018). As most of the large inversions were in the regions with low recombination rate in the genetic maps, it suggested that these inversions might be the orientation errors in the BTx623 reference genome. Similar observation was also reported from the published RTx430 genome (Deschamps et al., 2018).

**Figure 3.**
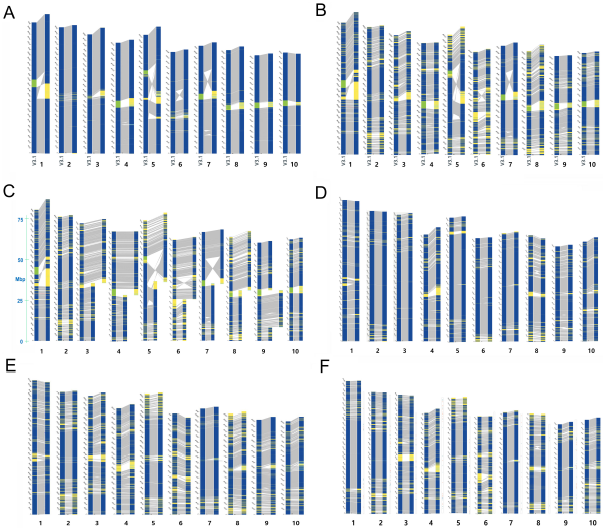
Alignments of Bionano maps between sorghum genomes. **(A)** BTx623 v3.1 reference sequence and BTx623 Bionano map; **(B)** BTx623 v3.1 reference sequence and RTx436 Bionano map; **(C)** BTx623 v3.1 reference sequence and Tx2783 Bionano map; **(D)** BTx623 Bionano map and Tx2783 platinum sequence; **(E)** BTx623 Bionano map and RTx436 platinum sequence; **(F)** Tx2783 platinum sequence and RTx436 Bionano map.

Based on this assessment, for the remaining analyses Tx2783 was used to characterize single-nucleotide variants (SNVs or SNPs) in the BTx623, Rio, RTx436, and RTx430 genomes (Fig. 4A). RTx430 had the most SNPs among the four genomes (Supplemental Table S5). The bulk of the SNPs, 92.6% on average, were predominantly called in intergenic regions; however, in gene regions, SNPs within the CDS regions of exons, which were the most highly conserved, were likely to have large effects (Fig. 4B). To identify large-effect SNPs within genes, we generated SIFT (sort intolerant from tolerant) scores using the Tx2783 genome annotation. The results revealed that 766, 1000, 1093, and 1004 genes in BTx623, Rio, RTx430, and RTx436, respectively, contained SNPs with SIFT scores◻<◻0.05 and were therefore predicted to be deleterious mutations. Among these genes, 183 genes were shared by all four genomes (Fig. 4C). Gene Ontology analysis revealed that these genes were associated with molecular function terms related to transferase activity, catalytic activity, and binding activity and biological process terms related to nitrogen compound metabolic process (Fig. 4D, Pearson Chi-Square test, P◻<◻0.05).

**Figure 4.**
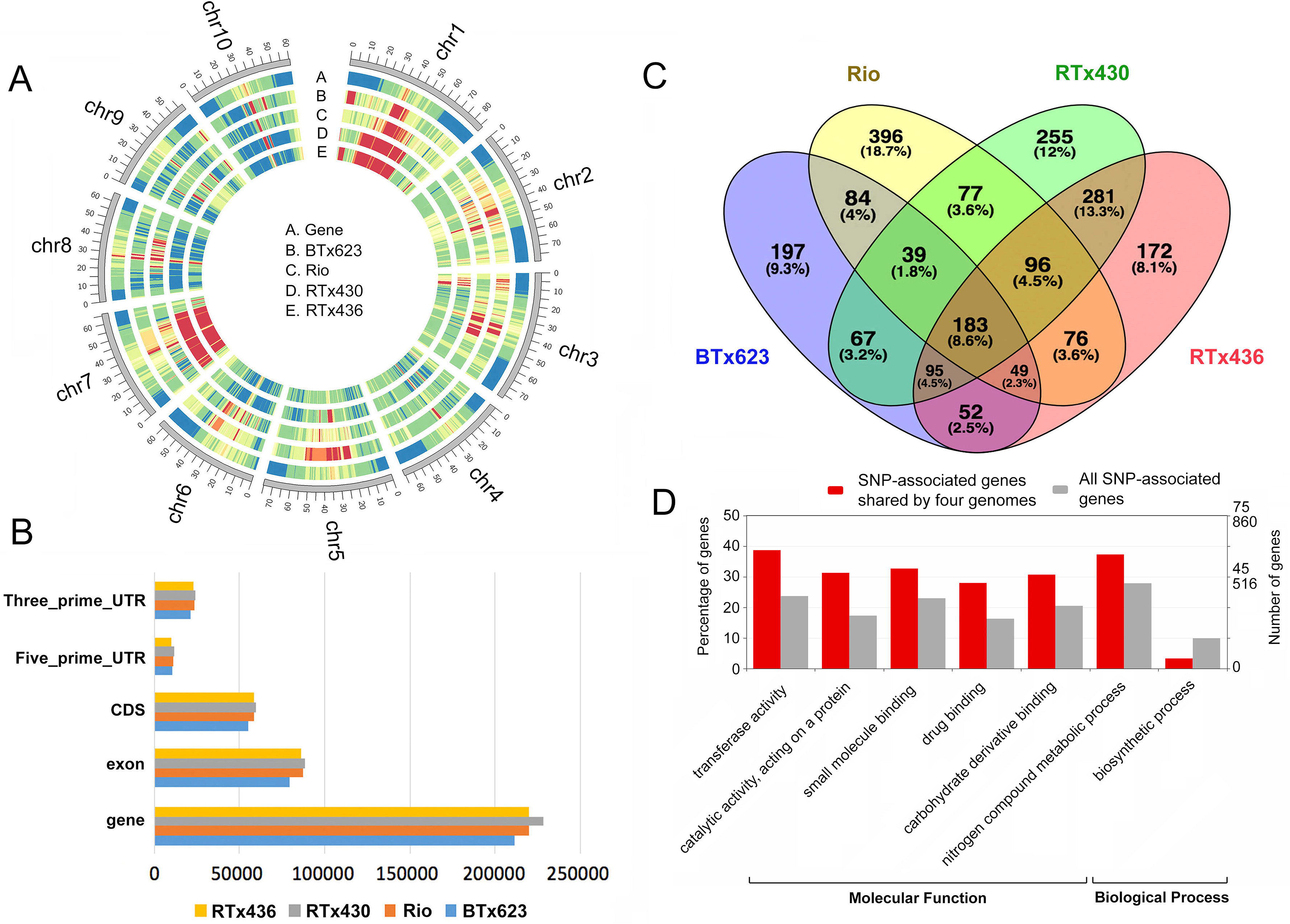
Characterization of SNPs in sorghum genomes. **(A)** Genome-wide distribution of SNPs in each genome relative to the Tx2783 reference. **(B)** Distribution of SNPs in the indicated gene features. **(C)** Venn diagram of deleterious mutations in each genome, using Tx2783 as a reference. **(D)** Gene Ontology enrichment of SNP-associated genes.

Although less common than SNPs, SVs have a greater potential to impact function due to their larger size and the possibility of altering gene structure, dosage, or location (Fuentes et al. 2019). We characterized SVs in BTx623, RTx436, RTx430, and Rio by whole-genome alignment using sorghum Tx2783 as the reference. Variants were classified into four different categories: insertion (INS), deletion (DEL), inversion (INV), and copy number variation (CNV). The largest number of SVs was identified in RTx430 (Figure 4A, Supplemental Table S6, Supplemental Data S2). Like the SNPs, these SVs were enriched in chromosome-terminal regions in all five genomes, but the specific SV categories that were enriched differed among individual genomes (Fig. 5A). In addition, we identified genotype-specific and common SVs in each genome (Fig. 5B to 5E), the majority of which were in intergenic regions; however, as noted above, SVs within genes were more likely to contribute to large-effect mutations (Fig. 5F–G). In addition, we also saw that the number of genes within SVs were correlated with the size of the SVs (Supplemental Fig. S1), and that overall, DEL SVs covered the most genes.

**Figure 5.**
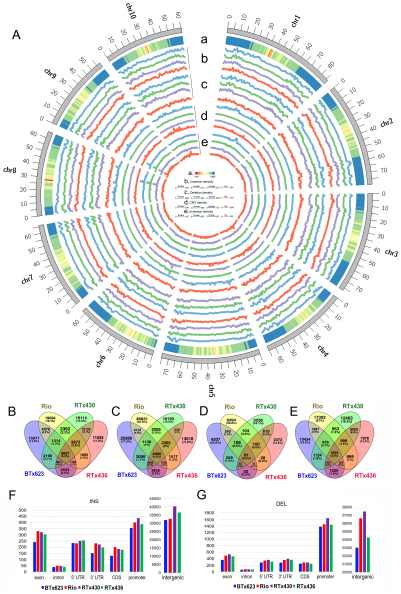
Characterization of large structural variations in sorghum genomes. **(A)** Circular visualization of different categories of SVs at the genome-wide level. a. Heatmap of gene density in the Tx2783 genome. b. Genome-wide distribution of INS density in four genomes: BTx623, RTx436, RTx430, and Rio. c. Genome-wide distribution of DEL density in four genomes: BTx623, RTx436, RTx430 and Rio. d. Genome-wide distribution of CNV density in four genomes: BTx623, RTx436, RTx430 and Rio. e. Genome-wide distribution of INV density in four genomes: BTx623, RTx436, RTx430 and Rio. **(B–E)** Overlap of SVs among sorghum genomes. **(F)** Distribution of INS SVs in different genomic regions. **(G)** Distribution of DEL SVs in different genomic regions.

Combining the information on SNPs and SVs, we found that the number of SNPs was positively correlated with the number of INS and DEL SVs in the same genomic region; this correlation was lower for CNVs and absent for INV SVs (Supplemental Fig. S2). CNVs are a specific class of SVs that play important roles in plant development (Soyk et al., 2019). We also investigated CNVs in these genomes, and found that the number and size of CNVs were distributed similarly in each genome, although they were slightly smaller and less abundant in the BTx623 genome (Supplemental Fig. S3A), and smaller CNVs were usually present in more copies than larger CNVs (Supplemental Fig. S3B, Supplemental Fig. S4).

We also investigated the possible functional roles of the genes affected by different types of SVs. Gene Ontology analysis using WEGO (Ye et al., 2018) revealed that genes affected by INS/DEL/CNVs were enriched for biological process terms related to metabolic and cellular processes. Enriched molecular functions included catalytic activity and binding (Supplemental Fig. S5).

### Core and Dispensable Genes Identification Using 62 Genomes Resequencing Data

To assess the core and dispensable genes, we constructed a pan-genome using the Tx2783 assembly and novel genomic regions represented by the SV insertions from the four other assemblies (Supplemental Data S2). The pan-genome had an effective genome length of ~820LMb and contained 35355 genes (29,612 protein coding + 4204 non-coding from TX2783 + 1539 [1517 protein coding and 22 non-coding] SVs annotated genes from Rio, RTx430, RTx436, and BTx623). Using publicly available resequencing data for 62 sorghum accessions (Supplemental Data S3; Mace et al., 2013; LeBauer et al., 2020; https://datacommons.cyverse.org/browse/iplant/home/shared/terraref), we aligned sequence reads were aligned to the sorghum pan-genome using BWA software (Li et al., 2009a). Mapping rates varied from 74.57% to 98.59%, averaging 93.25% in the resequencing lines (Supplemental Data S4), and were lowest in wild weedy lines (AusTRCF 317961, Greenleaf, PI330272, PI300119, Kilo and Zengada). The observed differences in mapping rates may be due to divergence between the sequenced genotypes. Using a conservative quality filter pipeline, we identified 8,052,666 SNPs throughout the genome in *S. bicolor* alone. SNPs were used to calculate the genetic distance between individuals, and a neighbour-joining tree was constructed using RAxML-NG (Kozlov et al., 2019) (Fig. 6A). In general, our phylogenetic tree was in consistent with a previous study by Mace et al.

Core and dispensable genes were identified by using genes covered in each of the 62 accessions under a 75% coverage cutoff. As a result, among the protein-coding genes, we identified 17,314 as core (present in all 62 lines), 3,466 as softcore (present in 59–61 lines), 9,952 as dispensable (present in 2–58 lines) and 173 as private genes (present in only one line) (Fig. 6B–C; Supplemental Table S7). Among the non-coding genes, we identified 1,803 as core, 734 as softcore, 1,906 as dispensable and 7 as private genes. (Supplemental Fig. S6, Supplemental Table S7).

**Figure 6.**
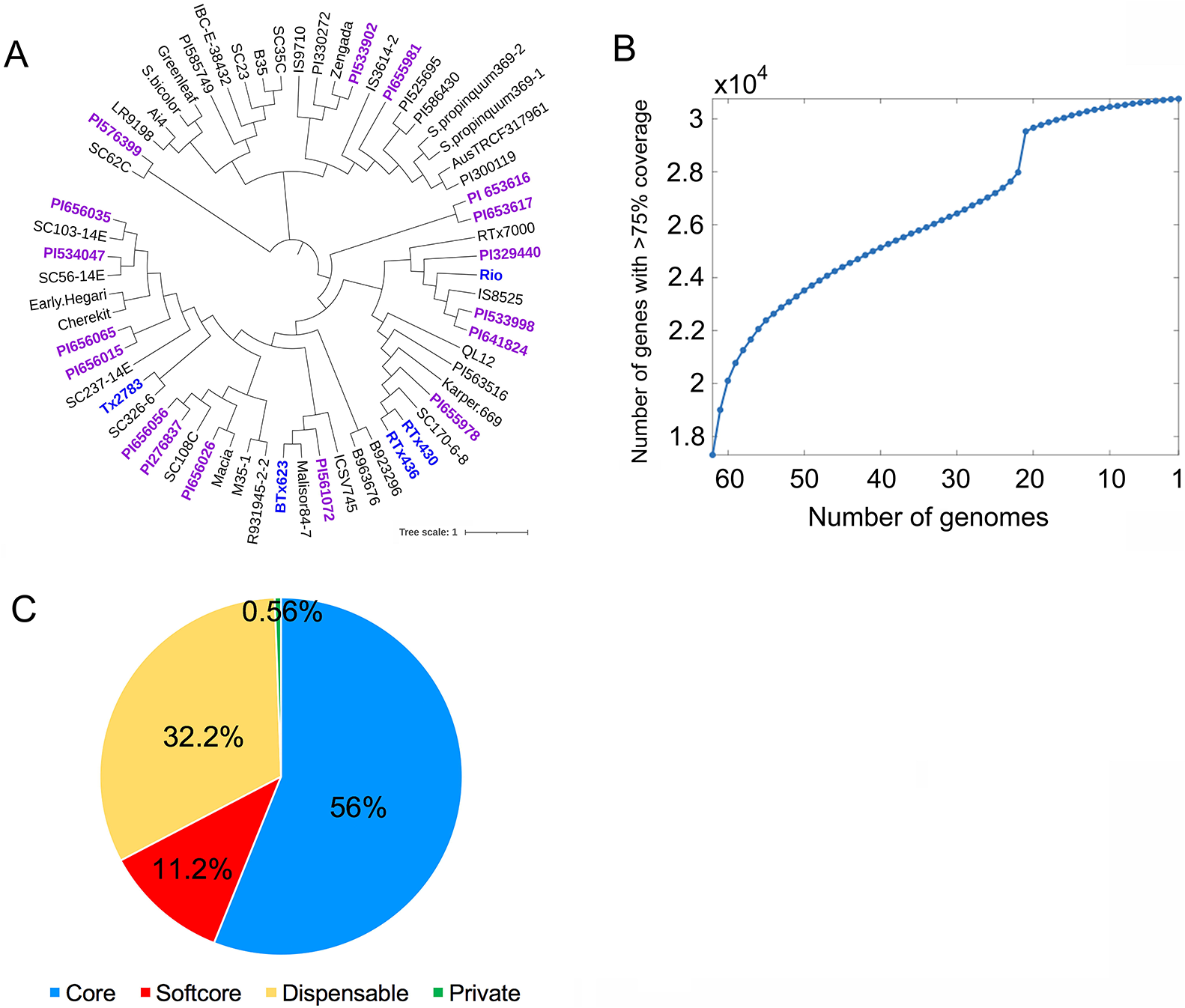
Phylogenetic tree of 62 sorghum accessions and distribution of core genes identified from the population. **(A)** Phylogenetic tree of 62 genome accessions. Blue: Five complete genome assemblies; Purple: 17 BAP/SAP lines; Black: Lines from Mace et al., 2013. **(B)** Cumulative curve of covered protein-coding genes in 62 genomes under 75% gene coverage cutoff. **(C)** Distribution of core, softcore, dispensable, and private protein-coding genes identified from 62 sorghum accessions using the 75% gene coverage cutoff.

### NBS-LRR Genes in Sorghum Genomes

To characterize disease resistance (R) genes, we first assessed the distribution of known NBS-LRR genes in both maize and sorghum (Supplemental Fig. S7). The results revealed that maize has far fewer R genes than sorghum (Paterson et al., 2009; Schnable et al., 2009). Whole-genome scanning of each sorghum line revealed enrichment of R genes on chromosomes 5 and 8 (Fig. 7A), with RTx430 having the most R genes (363) and Rio having the fewest (319) (Supplemental Table S8).

**Figure 7.**
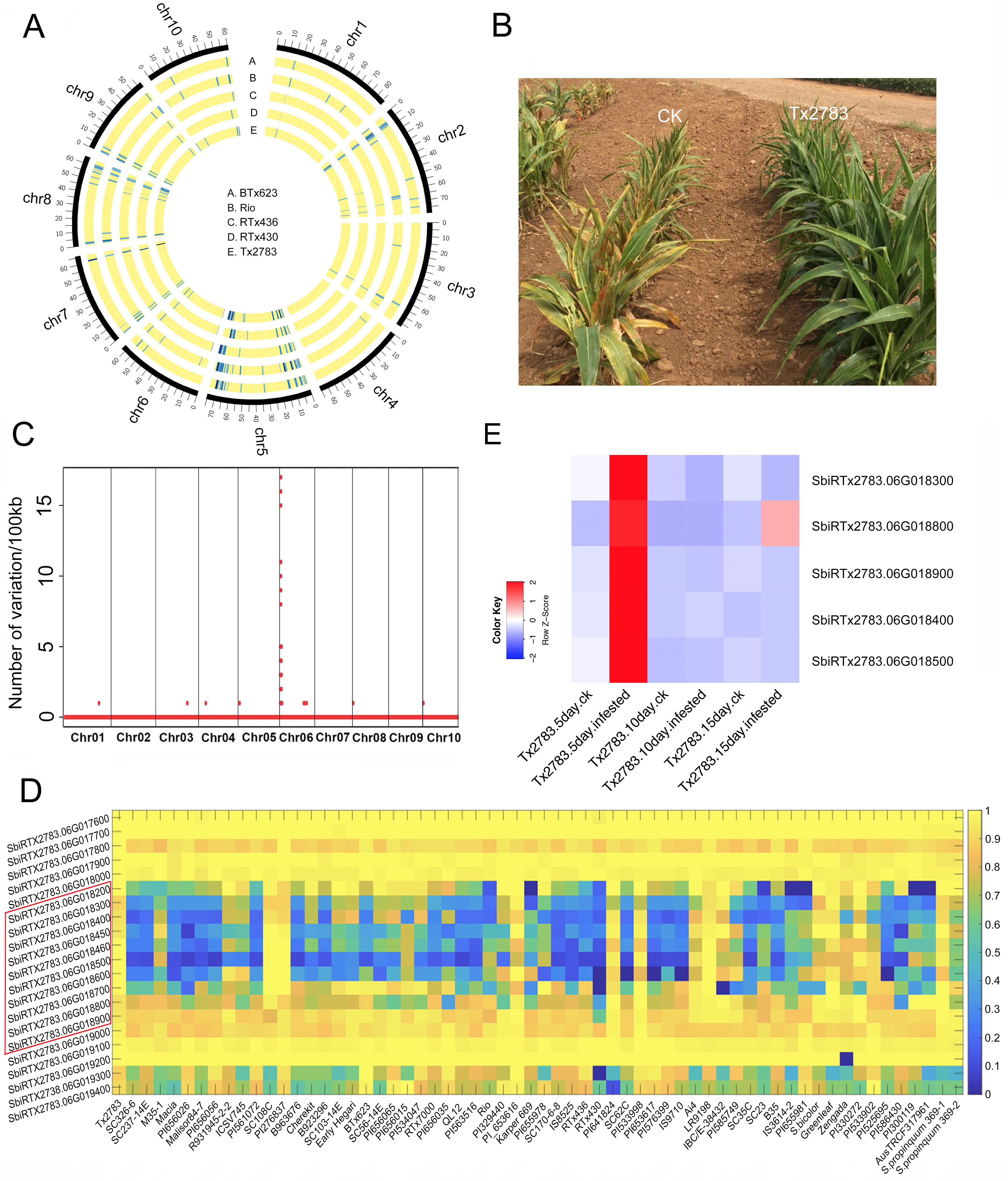
Identification and characterization of sugarcane aphid resistance locus. **(A)** Genome-wide distribution of NLR loci in each genome. **(B)** Phenotype of SCA resistance in sorghum line Tx2783. **(C)** Region of SCA resistance genes, identified using a RIL population. **(D)** Heatmap of genes within a 580-kb window of the SCA region across 62 sorghum accessions from resequencing data. Resistance genes are outlined in red on the left of the heatmap: SbiRTX2783.06G018200; SbiRTX2783.06G018300; SbiRTX2783.06G018400; SbiRTX2783.06G018450; SbiRTX2783.06G018460; SbiRTX2783.06G018500; SbiRTX2783.06G018600; SbiRTX2783.06G018700; SbiRTX2783.06G018800; SbiRTX2783.06G018900. **(E)** Expression of five genes in the SCA region from RNA-seq data. SbiRTX2783.06G018300; SbiRTX2783.06G018400; SbiRTX2783.06G018500; SbiRTX2783.06G018800; SbiRTX2783.06G018900.

In 2013, sugarcane aphid (SCA) emerged as a major insect pest of sorghum crops in North America (Alkan et al. 2011). The line Tx2783 is highly resistant to SCA, and thus has great potential to boost breeding targeted at aphid resistance (Tetreault et al., 2019) (Fig. 7B). Our genetic tests showed that the resistance allele in Tx2783 is dominant. To identify the resistance loci in Tx2783, we constructed a RIL population between Tx2783 and BTx623 and used it to generate two pools: one with 20 SCA-resistant individuals, and another with 20 susceptible individuals. After comparing the genotypes of the parental lines and the two RIL pools, we identified one peak on chromosome 6 harboring 17 large-effect variations as the candidate causal region in Tx2783 (Chr6: 3,008,428– 3,286,874) (Fig. 7C, Supplemental Data S5). Our SV analysis revealed 11 deletions in this region in the susceptible line BTx623; the largest deletion was 191,539 bp, and the remaining deletions were relatively small (< 3000 bp). (Supplemental Table S9, Supplemental Fig. S8). We then checked the overlapping genes in this region in the Tx2783 genome, and found that eight genes overlapped with three deletions in this region (Supplemental Table S9). Functional annotation of these genes showed that six of them encode proteins containing leucine-rich repeat (LRR) domains (Supplemental Data S6), which are involved in a variety of biological processes, including disease resistance and immune responses.

Genome sequence analysis in the sugarcane aphid resistance QTL region showed that there is a nearly identical 70,267-bp partial tandem duplication, which increased the complexity of this region (Supplemental Fig. S9) and required refinement of the loci (see Methods). Previous study has revealed that resistance genes are usually clustered within genomes (Stein et al., 2018). Therefore, we explored the depth of coverage of genes in the SV region in our population data (Fig. 7D). Based on the resequencing data, there was little to no coverage of this region in all but ten accessions for the eight genes. There were ten R genes in this region in Tx2783, but only three in BTx623. Therefore, we speculated that the gene dosage of these gene clusters in Tx2783 play a large role in SCA resistance.

To investigate how gene expression is impacted by SCA, we remapped the RNA-Seq data before and after sugarcane aphid infestation from a previous study (Tetreault et al., 2019) to our Tx2783 genome. Gene expression analysis revealed that five out of the ten R genes expressed before and after SCA infestation (Fig. 7E). We also observed diverse expression patterns for all genes throughout the genome (Fig. 8A). We then grouped all genes into 16 clusters based on their expression levels (Fig. 8B). This analysis revealed that 3,944 genes were upregulated relative to control plants after 5, 10, and 15 days of infestation. In particular, 173 NLR genes were expressed before or after SCA infestation, and 27 NLR genes were continuously upregulated after 5, 10, and 15 days of infestation. The overall expression of most NLR genes increased after 5 and 10 days of infestation, with a slight change at 15 days (Fig. 8C).

**Figure 8.**
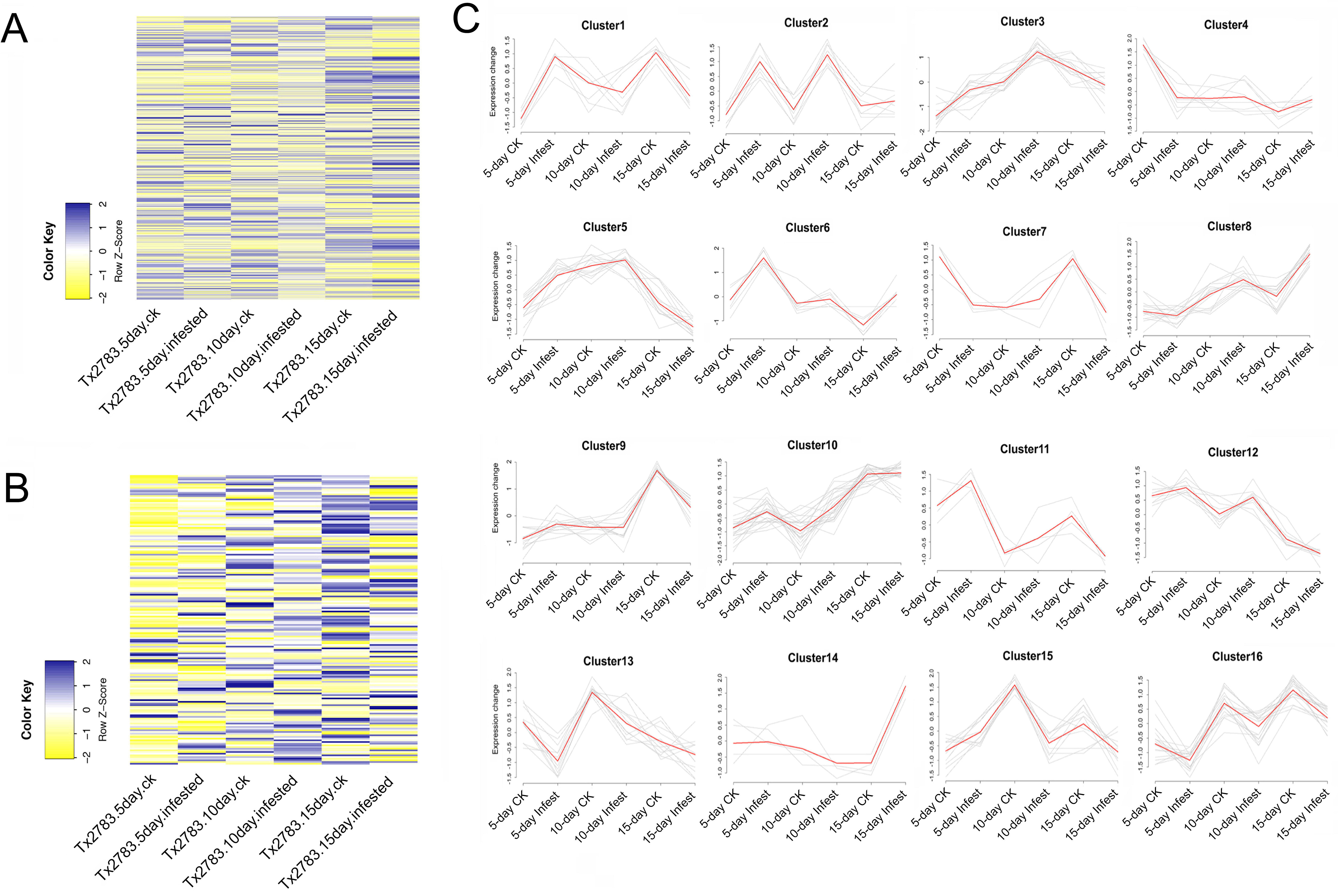
Gene expression analysis after sugarcane aphid infestation. **(A)** Heatmap of genes expressed before and after SCA infestation. **(B)** Clusters of genes expressed before and after SCA infestation. **(C)** Heatmap of NLR genes before and after SCA infestation.

## DISCUSSION

Sorghum is a very important cereal grain, forage, and bioenergy crop around the world. The first reference genome was released in 2009 (Paterson et al., 2009) and was updated 9 years later (McCormick et al., 2018). With the development of long-read sequencing technology, two other sorghum genomes have become available to the public: RTx430 (Deschamps et al., 2018) and Rio (Cooper et al., 2019). To date, however, no studies have examined SVs in sorghum genomes at the pan-genome level.

In this study, we generated two long-read sequenced genomes: Tx2783, which is highly SCA-resistant, and RTx436, which is a high-yielding, well-adapted pollinator line. Using these and other available genomes, we were able to investigate SVs in sorghum. Using Bionano maps, we found several incorrectly assembled regions in the current BTx623 reference genome; these assembly errors were confirmed by various map alignments. In light of this, we used the higher-quality Tx2783 genome as reference for calling SVs for subsequent analyses. The results revealed a large number of SVs in various sorghum genomes, and also they share some SVs at the same locus using the same reference; moreover, we found that different genomes can have SVs at different regions.

As the first examination of SVs across several genomes in sorghum, this study will provide valuable information to sorghum pan-genome comparisons and deeper insights into plant genomics. It is worth noting that even though genome quality has been dramatically improved by long-read sequencing technology, some complex regions, including recently tandem duplicated sequences, may still need to be refined because these regions may incorrectly assembled and collapsed, as the SCA region was in this study. Visual inspection of the hybrid scaffold at the SCA locus revealed an approximately 69.8-kb collapsed repeat in the contig assembly that was not resolved during the initial manual curation process. Similarly, alignment of the PacBio corrected reads to the contig assembly showed evidence of a collapsed assembly of approximately 77.7 kb within the QTL locus. To address this issue, we manually assembled the longest corrected reads from a 592-kb target region spanning the QTL region using GeneCodes Sequencher, yielding a new sequence that resolved the collapsed repeat such that it was congruent to the Bionano map and incorporated the previously soft-clipped reads.

SCA has become an aggressive pest of sorghum, causing severe yield losses. Therefore, identification of SCA resistance loci is urgently required and has great potential value. Tx2783 is a highly SCA-resistant line grown worldwide, and is thus an excellent resource for identification of resistance loci. In this study, we used a RIL population to map SCA resistance to a region on chromosome 6. We found several large insertions in this region in Tx2783 relative to the susceptible line BTx623. Eight genes overlapped with the SV region, of which six were in a 191-kb region, so it is reasonable to speculate that these genes or these SVs play important roles in SCA resistance. Our population analysis also showed that ten genes segregate with different sorghum accessions. We speculate that these ten gene clusters could function together in response to SCA as part of a complex resistance mechanism. Further studies in which one or all of these genes are mutated (e.g., using CRISPR technology) will be necessary to determine which genes confer the SCA-resistance phenotype, and to explore the underlying molecular mechanism. To broadly characterize R genes, we performed a whole-genome scan of NLR genes for each line. The results revealed that each sorghum genome has a similar number of R genes. Analysis of whole-genome distributions revealed that NLR genes are enriched on chromosomes 5 and 8; these R gene clusters could play important roles in resistance to SCA and other biotic stressors and diseases in sorghum. Our characterization of sorghum pan-genome variations and SCA resistance provides an important resource for the sorghum research community.

As mentioned above, genomics has entered the pan-genome era, in which the genetic diversity within a species is revealed through the availability of multiple genomes. On the other hand, the phenotypic differences among accessions are usually associated with dispensable and private genes. Consequently, it is important to distinguish the core and dispensable elements in a pan-genome. To construct a sorghum pan-genome, we used the Tx2783 genome as backbone and added insertions called from four other complete genomes. We then used genome resequencing data from 62 other sorghum accessions, including 44 accessions from Mace et al., 2013 and 17 additional BAP/SAP lines that add more diversity to the population, to identify the core and dispensable genes. After mapping these resequencing data to the pan-genome, SNPs were called and used to construct a phylogenetic tree, which was consistent with the findings of a previous study (Mace et al., 2013). We also observed that 21 lines had similar gene coverage among the population, resulting in a blip in the cumulative curve of covered genes. We then identified core and dispensable genes from the population. These different classes of genes are likely to explain the genetic and phenotypic diversification of sorghum species, providing fundamental information for population genomics in sorghum.

## METHODS

### Data Generation

#### Bionano Map

Ultra-high molecular weight nuclear DNA (uHMW nDNA) was isolated using a modified version of the Bionano Plant Tissue DNA Isolation Base Protocol (https://bionanogenomics.com/wp-content/uploads/2017/01/30068-Bionano-Prep-Plant-Tissue-DNA-Isolation-Protocol.pdf). Approximately 0.7–2 g of healthy young leaf tissue was collected from seedlings 2 weeks after germination. Leaf tissue was treated in a 2% formaldehyde fixing solution, washed, diced, and homogenized using a Qiagen TissueRuptor probe. The resultant homogenate was filtered through 100- and 40-μm cell strainers, and then pelleted by centrifugation at 2,500 × *g* for 20 minutes. Free nuclei were concentrated by step gradient centrifugation and pelleted by standard centrifugation at 2,500 × *g* for 10 minutes. The nuclei pellet was embedded into a low-melting-point agarose plug, followed by treatment with proteinase K and RNase A. Agarose plugs were washed four times in Bionano wash Buffer and five times in Tris-EDTA buffer, pH 8.0. Purified DNA was recovered by digesting the plug with agarase followed by drop dialysis against TE pH 8.0.

Direct Label and Stain (DLS) was used in combination with a Bionano Saphyr system to generate chromosome-level optical maps. Bionano DLS uses the DLE1 enzyme, which labels DNA molecules by recognizing CTTAAG sites and attaching a single fluorophore. DLS was performed using the Bionano Direct Label and Stain Kit (https://bionanogenomics.com/wp-content/uploads/2018/04/30206-Bionano-Prep-Direct-Label-and-Stain-DLS-Protocol.pdf) with a few modifications. Approximately 1 μg of sorghum uHMW nDNA was mixed with DLE-1 Enzyme, DL-Green label, and DLE-1 Buffer, and then incubated for 2:20 h at 37°C. The reaction was stopped by incubation for 20 min at 70°C, following by digestion with proteinase K for 1 h at 50 °C. Unincorporated DL-Green label was removed from the reaction by absorption onto a nitrocellulose membrane. The labeled, cleaned-up DNA sample was combined with Flow Buffer, DTT, incubated overnight at 4 °C, and quantified. The DNA backbone was stained by addition of Bionano DNA Stain at a final concentration of 1 microliter per 0.1 microgram of final DNA. Finally, the labeled, cleaned-up, and stained sample was loaded onto a single Bionano chip flow cell. DNA molecules were electrophoretically separated, stretched, imaged, and digitized using the Bionano Genomics Saphyr System and server (https://bionanogenomics.com/wp-content/uploads/2017/10/30143-Saphyr-System-User-Guide.pdf).

### PacBio and 10X Chromium Sequencing Collection

DNA was isolated from approximately 50 g fast-frozen young leaf tissue using a protocol described by Luo and Wing, 2003. Leaf material was ground in liquid nitrogen and incubated for 15 minutes in cold NIB buffer (10 mM Tris-HCl pH 8.0. 10 mM EDTA pH 8.0, 100 mM KCl, 500 mM sucrose, 4 mM spermidine trihydrochloride, 1 mM spermine tetrahydrochloride) containing 0.1% 2-mercaptoethanol. The resultant homogenate was filtered twice through Miracloth and mixed with 5% NIBT (NIB+10% Triton X-100). Free nuclei and cell debris were concentrated by centrifugation at 2,500 × *g* for 15 minutes and washed with NIB+0.1% 2-mercaptoethanol followed by a second centrifugation at 2,500 × *g* for 15 minutes. The pellet was embedded in LMP agarose plugs, and treated with proteinase K, melted, and digested with agarase. High-speed (30,000 × *g*) centrifugation cycles were performed to concentrate and remove solids and recover the DNA in the supernatant.

PacBio library preparation was performed using the Pacific Biosciences SMRTbell Template Prep Kit 1.0 following the protocol for > 30-kb libraries (https://www.pacb.com/wp-content/uploads/Procedure-Checklist-Preparing-HiFi-SMRTbell-Libraries-using-SMRTbell-Template-Prep-Kit-1.0.pdf) with some modifications. Prior to library construction, the DNA was sheared to approximately 50 kb using a BioRuptor (Diagenode) and pre-sized using the >20-kb high-pass broad-range protocol on a PippinHT (Sage Science). A second size selection for the final library was performed using the >20-kb broad-range PippinHT protocol, followed by a final DNA repair reaction. Quality of DNA at different steps of the process was evaluated using a FEMTO Pulse automated pulsed-field analyzer (Agilent Technologies, Wilmington, DE, USA). Quantification was performed using the Qubit dsDNA BR and HS reagents Assay kits on a Qubit 3.0 system. Sequencing was performed on a PacBio Sequel v6.0 system using 1M v3 SMRT Cells with 3.0 chemistry and diffusion loading. In total, six SMRT Cells each were used for Tx2783 and RTx436.

Chromium 10X was performed according to standard protocols using a non-sheared aliquot of the DNA used for PacBio library construction. Approximately 1 ng of DNA was loaded with 10X Chromium reagents and gel beads onto a Chromium chip (https://support.10xgenomics.com/de-novo-assembly/library-prep/doc/user-guide-chromium-genome-reagent-kit-v1-chemistry). The Chromium libraries were sequenced on an Illumina HiSeq2500 system. For Tx2783, a total of 231,083,110 clusters were obtained, corresponding to 69.8 Mbp or 87× genome coverage. For RTx436, a total of 229,285,050 clusters were obtained, corresponding to 69.3 Mbp or 86× genome coverage.

### Long- and Short-read Sequencing

Long-read data were generated using the Pacific BioSciences (Menlo Park, CA, USA) Sequel platform at Corteva Agriscience (Johnston, IA, USA). Six SMRT cells were sequenced for both samples with 10-hr movies and v6 chemistry. Raw subreads were filtered to a 3-kb minimum for both samples, generating 75× and 64× genome coverage for Tx2783 and RTx436, respectively. The raw subread N50 lengths were 23.1 kb (Tx2783) and 24.7 kb (RTx436).

Linked short-read data were generated by sequencing of 10X Genomics (Pleasanton, CA, USA) Chromium libraries at Corteva Agriscience on the HiSeq 2500 platform (Illumina, San Diego, California). The Tx2783 Chromium dataset had 76× coverage and mean molecule length of 62,942bp. The coverage depths and mean molecule lengths for the Tx2783 and RTx436 Chromium libraries were 76×/62.9kb and 84×/98.5 kb, respectively.

### Genome Assembly and Polishing

Raw PacBio subreads were corrected and assembled using Canu v1.8 (Koren et al., 2017) with the following parameters varying from the defaults: “correctedErrorRate=0.065 corMhapSensitivity=normal ovlMerDistinct=0.99”. The resultant contigs were filtered to a minimum contig length of 30 kb. Beyond the sequence consensus process that Canu performs after assembly, additional sequence polishing was performed by aligning raw PacBio subreads to the contig assembly using pbmm2 v0.12.0 and applying the Arrow algorithm from the Genomic Consensus package (v2.3.2) to get consensus calls. Both of these tools were obtained from pbbioconda (https://github.com/PacificBiosciences/pbbioconda). To further increase the consensus sequence accuracy, the long read contig assembly was complemented with Chromium linked short-read “clouds”, which have higher unique mappability than standard Illumina paired-end libraries. Chromium datasets were aligned to sequence contigs with Long Ranger v2.2.2. The assembly improvement tool Pilon v1.22 (https://github.com/broadinstitute/pilon) was used to correct individual base errors and small indels from the consensus of Chromium data aligned to the contigs using the parameter “--fix bases –minmq 30”. To decrease the compute time, a separate Pilon job was run for each contig against Chromium alignments specific to that contig. Contig-specific alignments were created using samtools v1.9 (https://github.com/samtools/samtools) from the Long Ranger output BAM file.

### Genome Mapping

Genome maps for Tx2783 and RTx436 were generating on the Bionano Genomics (San Diego, California) Saphyr platform at Corteva Agriscience using the Direct Label and Stain (DLS) system. For Tx2783, DLE-1–labeled molecule data from two flow cells (one chip) were filtered to create a data subset with a molecule N50 of 682 kb and 189× coverage. The filtered molecule dataset for Tx2783 was assembled via the Bionano Genomics Access software platform (Solve3.2.2_08222018) with the configuration file optArguments_nonhaplotype_noES_noCut_DLE1_saphyr.xml. The map assembly for Tx2783 consisted of 27 genome maps with a genome map N50 of 36.9 Mb and a total map length of 732.1 Mb.

For RTx436, DLE-1 labeled molecule data from two flow cells (1 chip) were filtered to create a dataset with a molecule N50 of 441 kb and 267× coverage. The filtered molecule dataset for RTx436 was assembled as described above, generating a map assembly of 28 genome maps with a genome map N50 of 37.7 Mb and a total map length of 723.6 Mb.

### Hybrid Scaffolding

An initial hybrid scaffolding was generated from the polished contigs and the Bionano genome maps using the Bionano Genomics Access software (Solve3.3_10252018) and the DLE-1 configuration file hybridScaffold_DLE1_config.xml to identify potentially problematic (smaller maps nearly identical to larger maps, low-coverage) or non-contributing genome maps. After this assessment, 19 genome maps from Tx2783 (Min length = 8.6 Mb) and 18 genome maps for RTx436 (Min length = 8.1 Mb) were selected to generate hybrid scaffolds. As part of the hybrid scaffolding process, chimeric mis-assemblies in the contigs were identified and cut to resolve conflicts relative to the genome maps. In addition to auto-conflict resolution, manual curation was performed to resolve overlapping contigs that were not addressed by the hybrid scaffolding workflow. A list of contig pairs that overlap in map space was generated from the gap file in the hybrid scaffold output directory, yielding 58 (Tx2783) and 51 (RTx436) overlapping pairs. Additional contigs cuts were added to the conflict resolution file to best resolve these issues. Embedded contigs are another assembly issue not resolved by the hybrid scaffolding workflow. A list of contig pairs in which a smaller contig is embedded within a collapsed region of a larger contig was also generated from the gap file in the hybrid scaffold output directory, yielding three (Tx2783) and eight (RTx436) embedded contig pairs. Similarly, additional conflict cuts were made in the larger contig to allow incorporation of the smaller contig after the hybrid scaffolding was re-run. In the final assembly, Tx2783 had 19 hybrid scaffolds (Scaffold N50= 36.0 Mb, Total scaffold length = 696.8 Mb) with 310 unscaffolded contigs with a total length of 27.1 Mb. RTx436 had 18 hybrid scaffolds (Scaffold N50= 37.6 Mb, Total scaffold length= 697.2 Mb) with 325 leftover contigs that were not scaffolded with a combined length of 25.0 Mb).

### Creating Pseudomolecules

With an average of 1.85 hybrid scaffolds per chromosome for the two lines, it is straightforward to create chromosome-scale pseudomolecules using the *Sorghum bicolor* v3.1 reference genome as a guide (https://phytozome.jgi.doe.gov/pz/portal.html#!info?alias=Org_Sbicolor). A generalized approach was used that rapidly maps scaffolds to a reference genome and determines their relative position and orientation. First, each scaffold was chunked into 100-bp fragments and then aligned to the *Sorghum bicolor* v3.1 reference genome using minimap2 v2.10 (Li, 2018). Then, a custom script was used to determine the chromosome position and orientation from the alignment of scaffold “read clouds”. All scaffolds were placed with this method. The remaining unscaffolded contigs were concatenated with 100-bp N-gaps and placed into Chr00.

### Integrating Plastid Genomes into the Assembly

It can be challenging to assemble plastid genomes from whole-genome PacBio libraries and distinguish them from plastid DNA integrated into the chromosome. An alternative approach is to insert a copy of an existing plastid genome reference sequence and to remove incomplete plastid assemblies from Chr00. A copy of the *Sorghum bicolor* chloroplast complete genome (GenBank: NC_008602.1) and *Sorghum bicolor* mitochondrion complete genome (GenBank: NC_008360.1) were added to the assembly as ChrC and ChrM, respectively. Only Tx2783 exhibited a SNP or small indel when the Canu-corrected reads were aligned with minimap2 and checked for variants with Pilon, which had a 1-bp deletion relative to the mitochondria reference. A modified version of ChrM incorporating this deletion was added to the TX2783 assembly. Incomplete plastid sequence contigs in Chr00 were identified by BLASTN (-F f -e 1e-200 -W32). Contigs with greater than 99% identity over more than 20 kb relative to the *Sorghum bicolor* plastid reference genomes were removed from the assembly. Manual curation of both assemblies was performed to fill gaps and better resolve the chromosomal chloroplast insertion on Chr09.

### Manual Curation of the SCA Locus

Visual inspection of the hybrid scaffold at the SCA locus revealed an approximately 69.8 kb collapsed repeat in the contig assembly that was not resolved during the initial manual curation process. In addition, alignment of the PacBio corrected reads to the contig assembly similarly showed evidence of a collapsed assembly of approximately 77.7 kb within the QTL locus (i.e. 2× corrected read coverage, soft-clipped reads flanking the potential repeat). The longest corrected reads from a 592-kb target region spanning the QTL region were manually assembled using GeneCodes Sequencher (Ann Arbor, Michigan) to generate a new sequence that resolved the collapsed repeat such that it was congruent to the Bionano map and incorporated the previously soft-clipped reads. As a result, 70,267 bp were added to the SCA locus. The entire manually curated region was swapped back into the pseudomolecule assembly and subjected to an additional round of sequence polishing with PacBio raw reads and 10X Chromium as described above. This was necessary because the corrected reads were used for the targeted assembly. Finally, the curated and polished 592-kb region was swapped back into the original pseudomolecule assembly. Thus, the final assembly, including the manually curated SCA locus was only subjected to a single round of polishing.

### Annotation of the Tx2783 and RTx436 Assembly

Gene annotations were performed using a strategy that combined evidence-based and *ab initio* gene predictions. For evidence-based predictions, genome-guided transcript assemblies were generated from four different assemblers, Trinity (v2.8.4) (Grabherr et al., 2011), StringTie (v1.3.5) (Pertea et al., 2015), Cufflinks (v2.1.1) (Trapnell et al., 2010), and PsiCLASS (v1.0.1) (Song et al., 2019), using the transcriptome data from seven tissue types across the juvenile, vegetative, and reproductive stages of development (Supplemental Data S7). The best sets of transcripts were identified and annotated as genes using Mikado (v2.0rc6) (Venturini et al., 2018). Briefly, the RNA-seq reads from each library were mapped to STAR (v2.7.0d) (Dobin et al., 2013)-indexed Tx2783 and RTx436 genomes using a two-pass mapping approach. Default options were used for mapping with few post-processing options enabled. Individually mapped RNA-seq libraries were then pooled, sorted, and indexed using Picard (v2.7.0) (http://broadinstitute.github.io/picard) for transcript assembly programs. For all genome-guided transcriptome assemblers, default options were used; if RNA-seq strandedness was allowed, it was set to stranded. For Trinity, maximum intron size was set to 10000. All assemblers generated a GFF3 as the final output except for Trinity, for which assemblies were mapped against the genome using GMAP (v2019-03-15) (Wu et al., 2005). Portcullis (v1.1.2) (Mapleson et al., 2018) was used to generate high-quality splice junctions from the merged mapped reads. Mikado was configured to use all transcript assemblies, portcullis-generated splice sites, and a plants.yaml scoring matrix. Preliminary transcripts were prepared for Mikado by 1) merging all transcripts and removing the redundant copies, 2) processed using TransDecoder (v5.5.0) (Grabherr et al., 2011) (to identify open reading frames), and 3) aligned with BLASTX (v2.2.29+) (Altschul et al., 1997) against SwissProt Viridiplantae proteins (to identify full-length transcripts). Default options were used for TransDecoder. For BLASTX, maximum target sequences were set to 5, and output format to xml. These were provided as input for Mikado for picking and annotating the best transcripts for each locus. The resultant GFF3 file was used to extract transcripts and proteins using the gffread utility from the Cufflinks package.

Additional structural improvements to Mikado-generated transcripts were completed using the PASA (v2.3.3) (Haas et al., 2003) genome annotation tool. The inputs for PASA included 209,835 maize EST derived from GenBank, Mikado-assembled transcripts for Tx2783 and RTx436, 37,655 sorghum iso-seq transcripts from 11 developmental tissues, and 31,881 sorghum full-length cDNAs from RIKEN that were filtered for intron retention (Reddy et al., 2016; Wang et al., 2018). PASA was run with default options; in the first step, transcript evidence was aligned to the masked sorghum genomes using GMAP (v2019-03-15) (Wu et al., 2005) and Blat (v.36) (Kent, 2002). The full-length cDNA and Iso-seq transcript IDs were passed in a text file (-f FL.acc.list) during the PASA alignment step. Valid near-perfect alignments with 95% identity were clustered based on genome mapping location and assembled into gene structures that included the maximal number of compatible transcript alignments. PASA assemblies were then compared with sorghum Mikado transcript models using default parameters. PASA updated the models, providing UTR extensions, as well as novel and additional alternative isoforms. PASA-generated models were passed through the MAKER-P (v3.0) (Campbell et al., 2014) annotation pipeline as model_gff along with all the transcript and protein sequences to yield Annotation Edit Distance (AED) scores to assess the quality of annotations. Transposon element (TE)-related genes were filtered using the TEsorter tool (Zhang et al., 2019), which uses the REXdb (viridiplantae_v3.0 + metazoa_v3) (Neumann et al., 2019) database of TEs. To supplement the evidence annotation, the protein sequences from Mikado transcripts and RNA-seq data were used for *ab initio* gene model prediction with BRAKER (Hoff et al., 2015). Non-overlapping BRAKER gene models were updated with PASA, filtered for TE-related genes, provided with AED scores using MAKER, and added to the evidence set. We further filtered this combined set with the criterion AED < 0.75, and then applied phylogeny filters by aligning protein sequences to maize, rice, *Brachypodium*, and Arabidopsis proteins to identify conserved and lineage-specific genes. The genes were loaded in Ensembl core databases and quality-checked to identify transcripts with incomplete CDSs, which were programmatically corrected and also checked for translation errors. Transcripts with complete CDSs were tagged as protein-coding and those with incomplete CDSs as non-coding. Functional annotation was performed using InterProscan-5.38-76.0 (Jones et al., 2014). Canonical assignment of transcripts was performed using a combination of criteria including domain coverage, translation length, and AED scores comparing each transcript to sample specific transcripts assembled with Stringtie. Transcripts were ranked by each criterion, and a canonical transcript was selected by a ranked-choice voting scheme (Olson et al., 2020). The functional and canonical attributes were added to the annotation files, and quality checking was performed using GFF3toolkit (Chen et al., 2019).

### Rampage Library Construction, Data Generation and Analysis

This protocol is a modified version of a previously published method (Batut et al., 2013). Prior to incubation with Terminator™ 5′-Phosphate-Dependent Exonuclease (TEX) (Lucigen) to remove all residual RNAs containing 5’ monophosphate, as described in Batut et al,. we removed all ribosomal RNAs using the RiboMinus™ Plant Kit for RNA-Seq (Thermo Fisher Scientific).

We then performed first-strand synthesis using the SMARTer Stranded Total RNA Kit V2-Pico Input Mammalian (Takara). Following purification with RNAclean XP (Beckman Coulter), 5’ cap oxidation, 5’ cap biotinylation, RNase I digestion, and streptavidin pulldown (Cap Trapping) were performed as described in Batut et al.

Amplification of purified cDNAs (two rounds of PCR to attach Illumina adapters and amplify the libraries) followed by AMPure XP cleanup (Beckman Coulter) was done using the “SMARTer Stranded Total RNA Kit v2” (Takara) according to protocol. All samples were processed separately, quantitated on a 2100 Bioanalyzer using a HS-DNA-Chip (Agilent), and adjusted to a concentration of 10 nM. Libraries were then pooled at equimolar concentration and sequenced on an Illumina NextSeq 550 Sequencer.

RAMPAGE reads were aligned to the reference using STAR 2.7 on SciApps (Wang et al., 2018). The mapped reads from each tissue were clustered using Paraclu (Frith et al., 2008). The BAM alignment files from STAR were converted to BED using the bamtobed tool and grouped using the groupBy tool from BEDTools v2.29.2 (Aaron et al., 2010) to sum up reads that started at the same position and on the same strand as the input to Paraclu. Paraclu was run with default settings with the minimal number of reads to form a cluster; -minValue was set to 10. The paraclu-cut.sh script within Paraclu was used to simplify and remove clusters with length > 200, max density/min density < 2, or that were contained within another cluster. Peaks with the clusters were identified using scripts provided in this Github repository: https://github.com/davetang/paraclu_prep. TSS profile plots for scores over the TSS region using an annotated 5’ UTR were generated using deeptools2 (Ramirez et al., 2016).

### SV Detection

All available genomes were aligned to the sorghum Tx2783 genome using MUMmer 4 (Marcais et al., 2018) with default parameters. Plots of the alignment only used the one-to-one alignment. SNP and small indels (<50 bp) were also called by the tool “show-snp” in the MUMmer 4 package. Large indels and inversions were called using the SVMU pipeline (Chakraborty et al., 2019). Insertions and deletions (>50 bp) with any overlap with the gap region in the reference genome were filtered out. Functional annotation of SNPs was performed using SnpEff (Cingolani et al., 2012) based on the Tx2783 genome annotation. SIFT 4G (Vaser et al., 2016) was used to predict deleterious mutations through the SciApps platform (Wang et al., 2018).

### Pan-Genome Construction and Resequencing Data Analysis

The pan-genome was constructed using Tx2783 as backbone, with the addition of the sequence of insertions which were called from the other four complete genomes. Resequencing data from Mace et al. 2013 and 17 BAP/SAP lines (https://datacommons.cyverse.org/browse/iplant/home/shared/terraref) were used for population analysis. Three lines (KS115; Yik.solate; Kilo) were removed due to low gene coverage; the remaining 61 genomes were mapped to this pan-genome using the BWA software (Li et al., 2009a). SAMtools (Li et al., 2009b) was used to convert mapping results to bam format. On one hand, BEDTools v2.29.2 (Aaron et al., 2010) was used to convert the bam files to bed format and calculate the gene coverage from the reads across 62 genomes. Next, core, softcore, dispensable and private genes were identified under three different cutoffs based on coverage: 50%; 75% and 90%; On the other hand, bam alignments were converted to pileup format using the pileup command. SNPs were called using both minimum base quality threshold and minimum mapping quality of 20. We then used vcf2phylip v2.0 (Ortiz, 2019) to convert the VCF to Phylip format, and then the python script ascbias.py (https://github.com/btmartin721/raxml_ascbias) was used to remove sites considered as invariable by RAxML. Finally, RAxML-NG (Kozlov et al., 2019) was used to estimate the best ML tree, then the bootstraps were used to plot support on the best tree.

### Identification of NLR Loci and Gene Expression Analysis

Genome-wide nucleotide-binding and leucine-rich repeat (NLR) loci were annotated using NLR-Annotator (Steuernagel et al., 2020), and loci that overlapped with genes were extracted using BEDTools (Quinlan et al., 2010). Phylogenetic trees were constructed using the Interactive Tree Of Life (iTOL) (Letunic et al., 2019). Illumina raw reads from Tetreault et al (Tetreault et al., 2019) were aligned to the Tx2783 reference genome using STAR with a minimum intron length set to 20Lbp and a maximum intron length set to 50Lkb, with default settings for other parameters. Quantification of genes and isoforms was performed using cufflinks version 2.2.1. A k-means approach was used [R Bioconductor package ‘Mfuzz’ (Kumar et al., 2007)] to cluster dynamically expressed genes based on their expression profiles across different time points after sugarcane aphid infestation.

### Identification of the SCA Resistance Locus

To identify the SCA resistance locus, the two parental lines Tx2783 and BTx623 and two pools from a recombinant inbred line (RIL) population, one consisting of 20 SCA-resistant individuals and the other consisting of 20 SCA-sensitive individuals, were sequenced on the Illumina platform. The average sequencing depth was about 10×. Bowtie2 (Langmead et al., 2012) was used to align the reads to the sorghum BTx623 v3.1.1 reference genome in ‘--very-sensitive’ mode. Duplicates from sequencing were masked using Picard tools (http://broadinstitute.github.io/picard/). After alignment, variations in the four samples were called by samtools (Li et al., 2009), using only reads with sequencing and alignment quality above 20. Only sites sequenced in all four samples were considered for the subsequent analysis. Candidate variations were required to meet the following criteria: 1) homozygous in all four samples; 2) different genotypes in the two parental lines; 3) BTx623 and Tx2783 genotype in the sensitive and resistant pools, respectively. The effects of the remaining variants were then predicted by SNPeff (Cingolani et al., 2012). Only variations annotated with MISSENSE, NONSENSE, or SPLICE_SITE_ACCEPTOR/DONOR effects were considered to be large-effect variations. The variations remaining as candidates were counted in 100-kb sliding windows with a step size of 10 kb.

## ACCESSION NUMBERS

The PacBio and Bionano data of the sorghum Tx2783 and RTx436 genomes generated in this study have been deposited in the European Nucleotide Archive (ENA) under accession numbers ERS4546874 and ERS4546867, respectively. 10X Chromium sequencing data are available under accession ERS4804287 and ERS4804288 for Tx2783 and RTx436 respectively. Genome assemblies of Tx2783 and RTx436 are available under accession numbers ERZ1345851 and ERZ1345852, respectively. RIL population sequencing data are available under accession number PRJEB39196. RAMPAGE datasets are available under accession number PRJEB42222.

---

**Table 1. Genome assembly statistics for Tx2783 and RTx436.**

## SUPPLEMENTAL INFORMATION

**Supplemental Fig. S1** Number of different types of SVs and the number of overlapping genes in the indicated genomes: (**A–B**). BTx623; (**C–D**). Rio genome; (**E–F**). RTx436 genome; (**G–H**). RTx430 genome.

**Supplemental Fig. S2** Correlation of number of different SVs with the number of SNPs in each sorghum genome.

**Supplemental Fig. S3** (**A**). Distribution of number of CNVs per gene in each genome. (**B**). Distribution of size of CNVs in each genome.

**Supplemental Fig. S4** Correlation of number of CNVs with Size of CNVs in each sorghum genome.

**Supplemental Fig. S5** Gene Ontology enrichment of genes overlapping various SVs: (**A**) Insertion. (**B**) Deletion. (**C**). CNV.

**Supplemental Fig. S6 Characterization of core non-coding genes.**(A) Cumulative curve of non-coding genes across 62 sorghum accessions. (B) Distribution of core, softcore, dispensable and private non-coding genes identified from 62 sorghum accessions.

**Supplemental Fig. S7** Chromosome distribution of NBS-LRR genes in sorghum (**A**) and maize (**B**).

**Supplemental Fig. S8** Manual curation of assembly in the SCA region.

**Supplemental Fig. S9** Dotplot of deletions in the SCA resistance region between Tx2783 and BTx623.

**Supplemental Table S1.** Summary of Bionano data for Tx2783, RTx436 and BTx623.

**Supplemental Table S2.** Summary of TE annotations for Tx2783, RTx436, and BTx623 genomes from EDTA.

**Supplemental Table S3.** Status of protein-coding gene annotations for Tx2783 and RTx436.

**Supplemental Table S4.** The list of large inversions that detected in Tx2783 and RTx436.

**Supplemental Table S5.** Number of SNPs called in each genome relative to the Tx2783 reference.

**Supplemental Table S6.** Number of SVs called in each genome relative to the Tx2783 reference.

**Supplemental Table S7.** Number of core, softcore, dispensable and private genes under various cutoffs of gene coverage for protein-coding and non-coding genes identified from 62 sorghum accessions.

**Supplemental Table S8.** Number of NLR loci in each genome, identified by genome scanning.

**Supplemental Table S9.** Summary of SVs in the SCA resistance region.

**Supplemental Data S1.** Structural variations called from Bionano genome maps between different genomes.

**Supplemental Data S2.** Structural variations between different genomes called from whole-genome sequence alignments using Tx2783 as reference.

**Supplemental Data S3.** Information of germplasm that was used for this study.

**Supplemental Data S4.** Mapping status of different sorghum accessions to the pan genome.

**Supplemental Data S5.** Large-effect SNPs in BTx623 and Tx2783 at SCA region.

**Supplemental Data S6.** InterProScan result of genes within the SCA region.

**Supplemental Data S7.** RNA-seq data that is used for genome annotation.

## AUTHOR CONTRIBUTIONS

W., Y.J., and D.W. conceived the idea for the study; V.L and K.F. generated genome assemblies; K.C., A.O. B.W. XH.W., Y.J. and XF.W generated genome annotations; M.R., T.G, JD. generated RNA, RNA-seq and RAMPAGE libraries; B.W analyzed the SNP and SV: B.W, L.W and A.O population analyses and core and dispensable gene. C.H. and Z.X. provided SCA seeds; S.A. performed the test of Peterson’s sorghum lines for sugarcane aphid resistance; J.H. developed and selected the SCA-resistance RIL population. Y.H. constructed the sugarcane aphid resistance RIL population; Y.J. mapped the sugarcane aphid resistance region; B.W., Y.J., and D.W. wrote the manuscript.

## ACKNOWLEDGEMENTS

We thank John Burke, Geoffrey Morris, Bob Klein, Bill Rooney, and Gary Peterson for helpful input concerning germplasm selection. This work was supported by USDA-SCA grant #51530311 and NSF Gramene grant #52930511. The authors thank Peter Van Buren for help with the computational system.

