## Supplemental Table for "Pan-genome Analysis in Sorghum Highlights the Extent of Genomic Variation and Sugarcane Aphid Resistance Genes": Supplemental Information.docx

**Supplemental Figures**

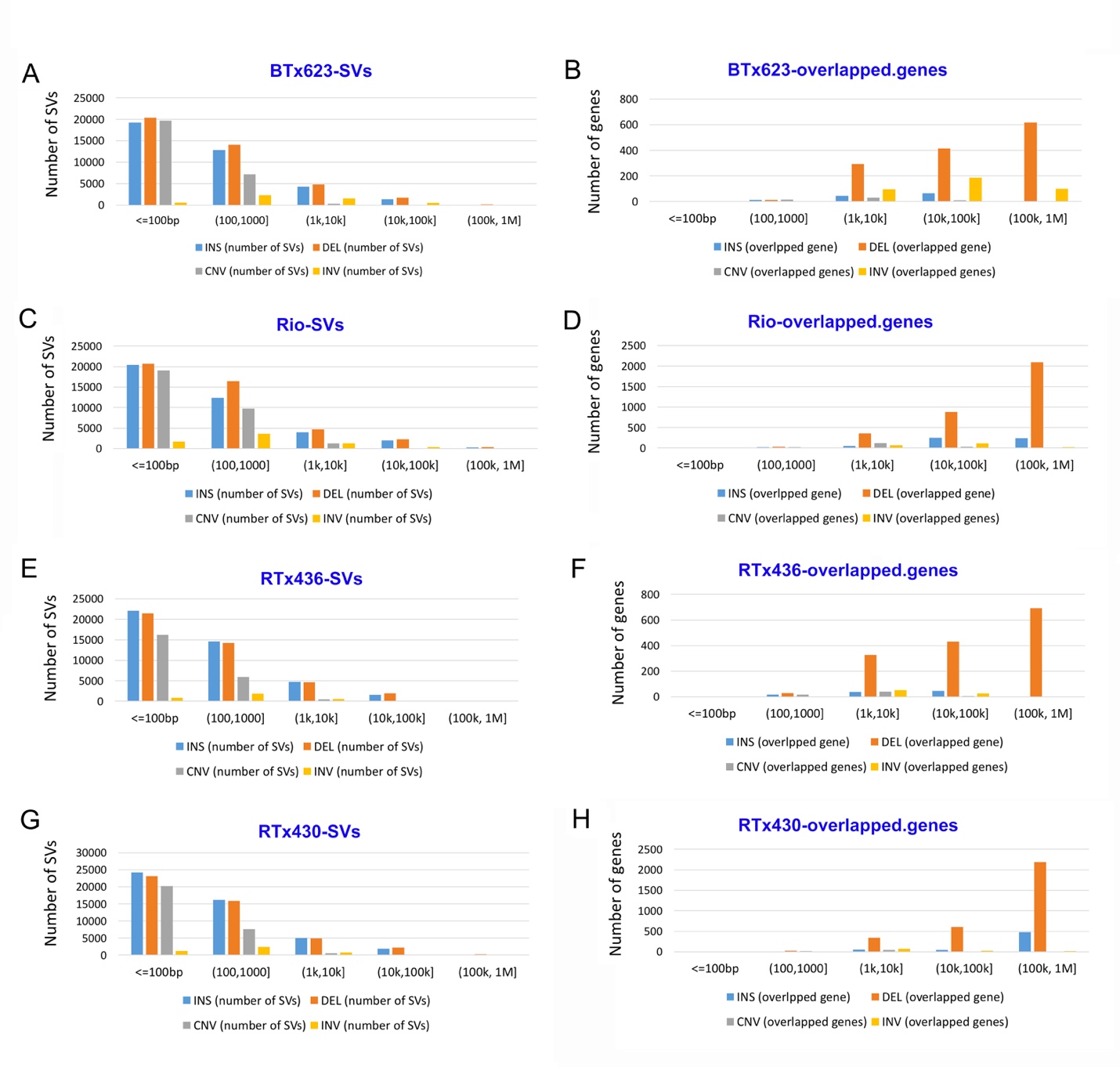

**Supplemental Fig. S1** Number of different types of SVs and the number of overlapping genes in the indicated genomes: (**A–B**) BTx623; (**C–D**) Rio genome; (**E–F**) RTx436 genome; (**G–H**) RTx430 genome.

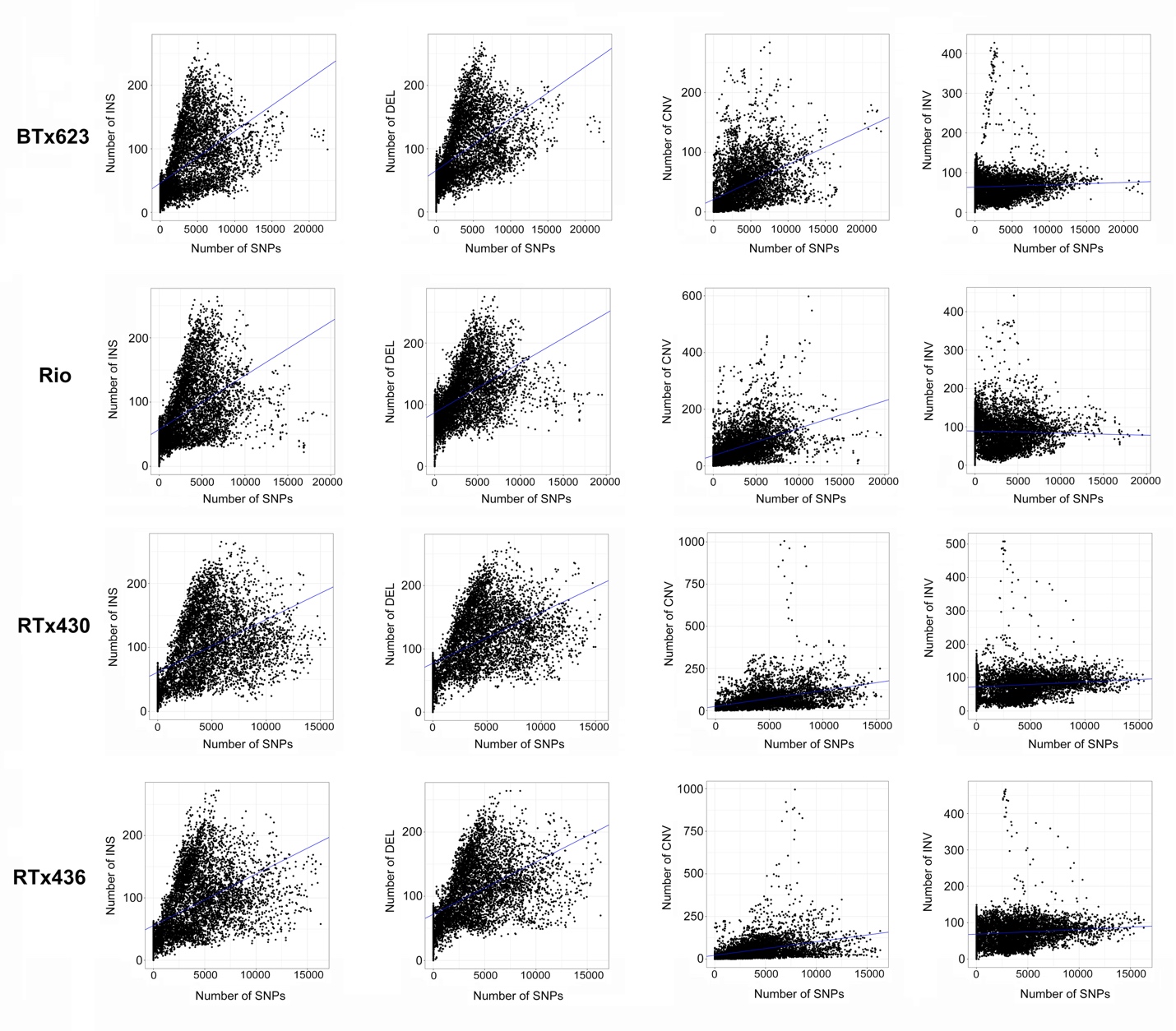

**Supplemental Fig. S2** Correlation of number of different SVs with the number of SNPs in each sorghum genome.

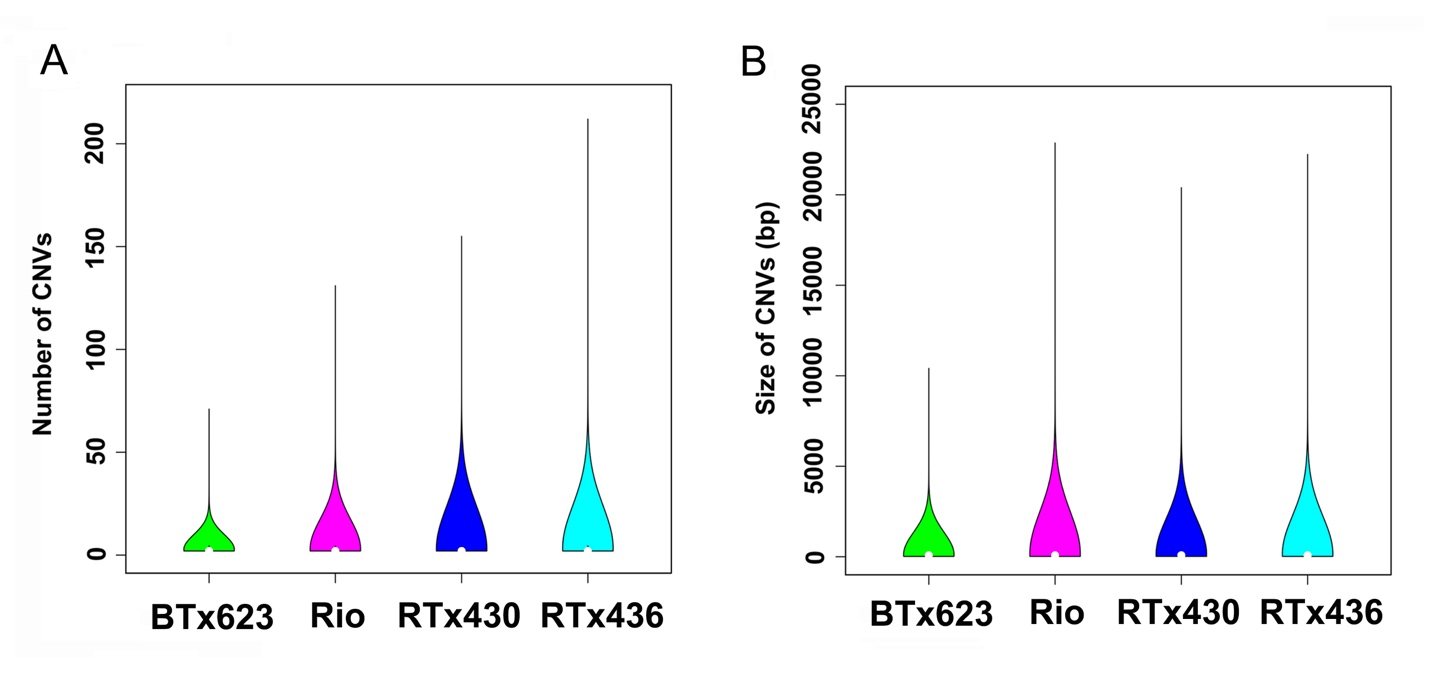

**Supplemental Fig. S3** (**A**) Distribution of number of CNVs per gene in each genome. (**B**) Distribution of size of CNVs in each genome.

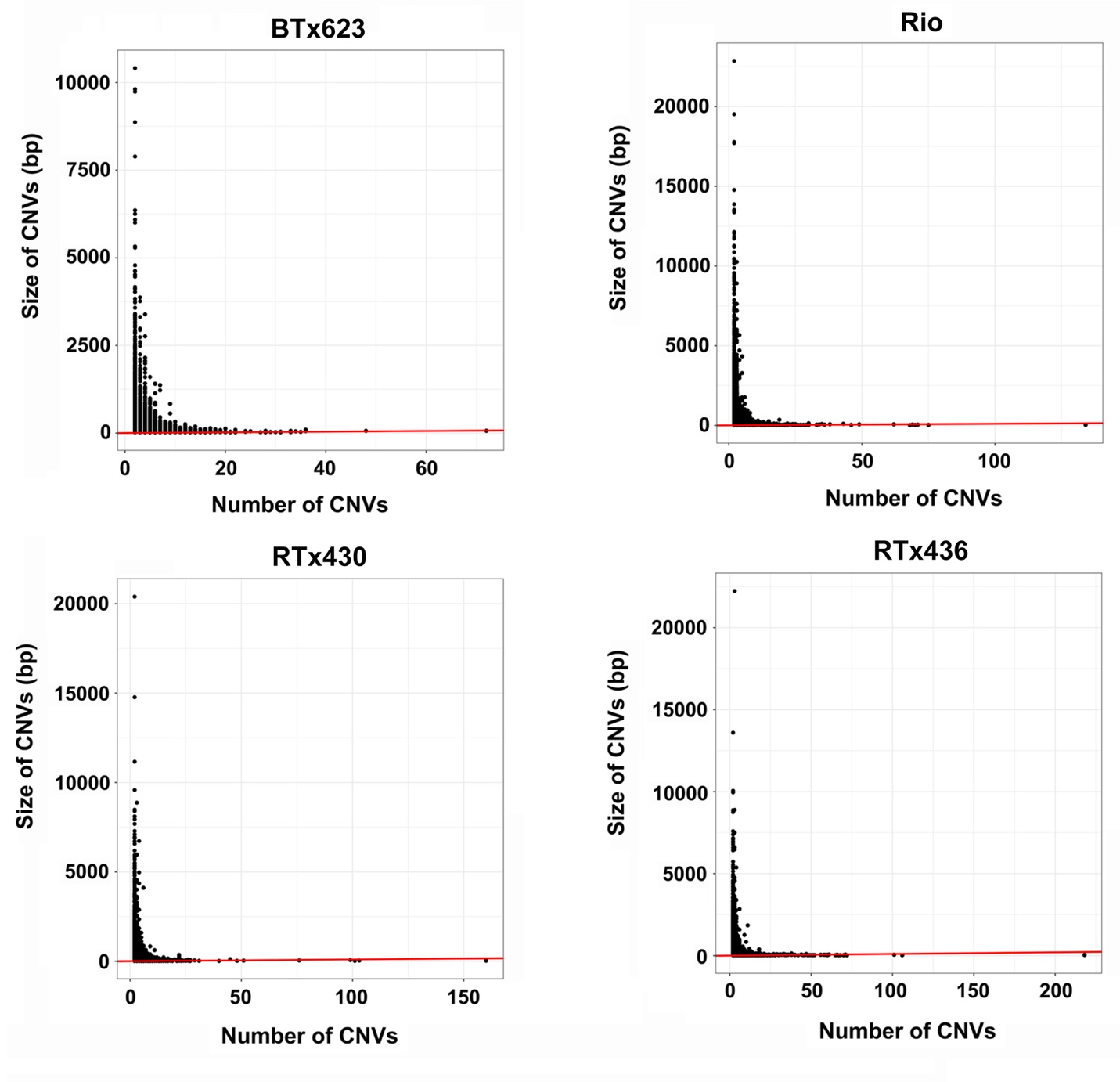

**Supplemental Fig. S4** Correlation of number of CNVs with size of CNVs in each sorghum genome.

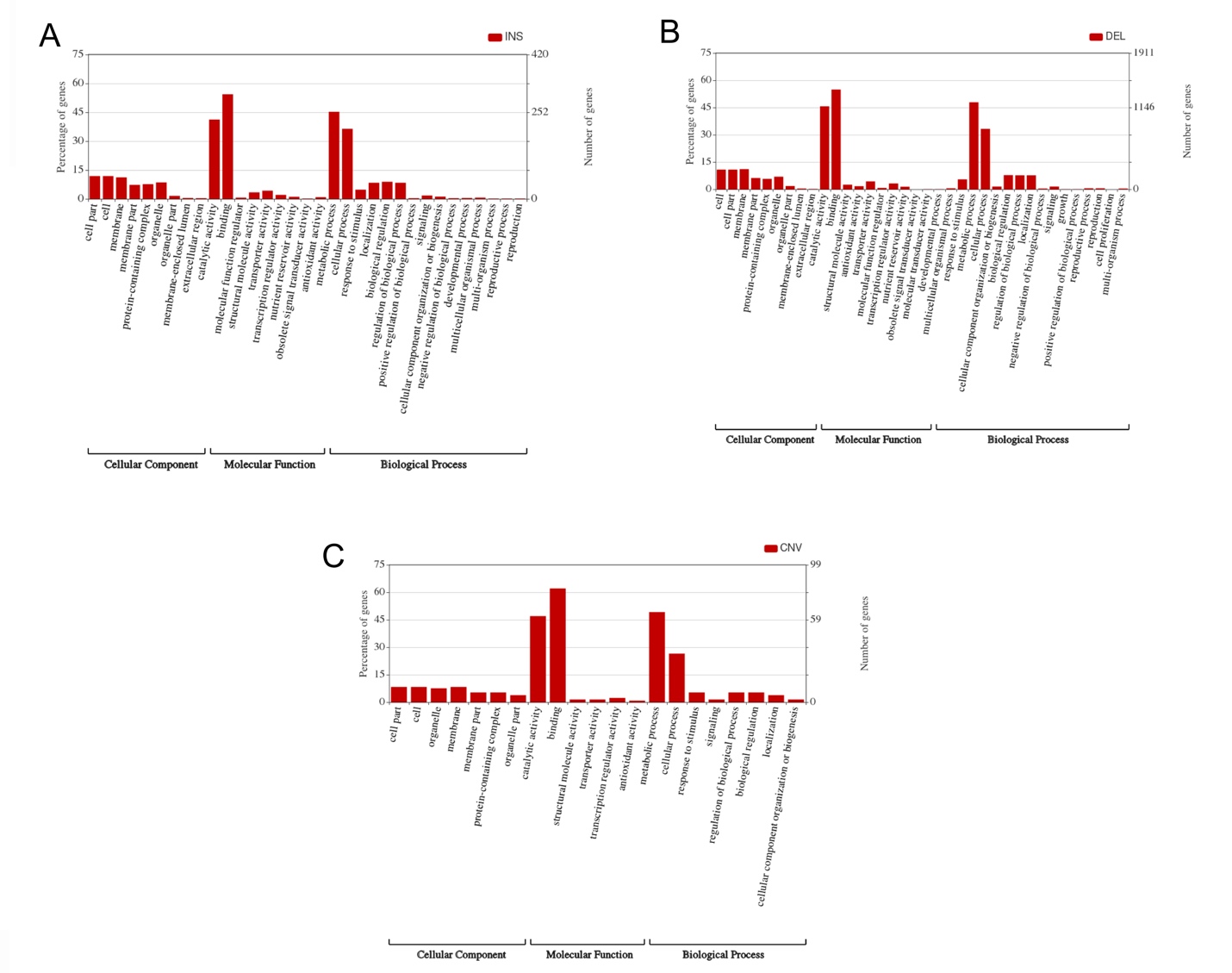

**Supplemental Fig. S5** Gene Ontology enrichment of genes overlapped with different SVs: (**A**) Insertion. (**B**) Deletion. (**C**) CNV.

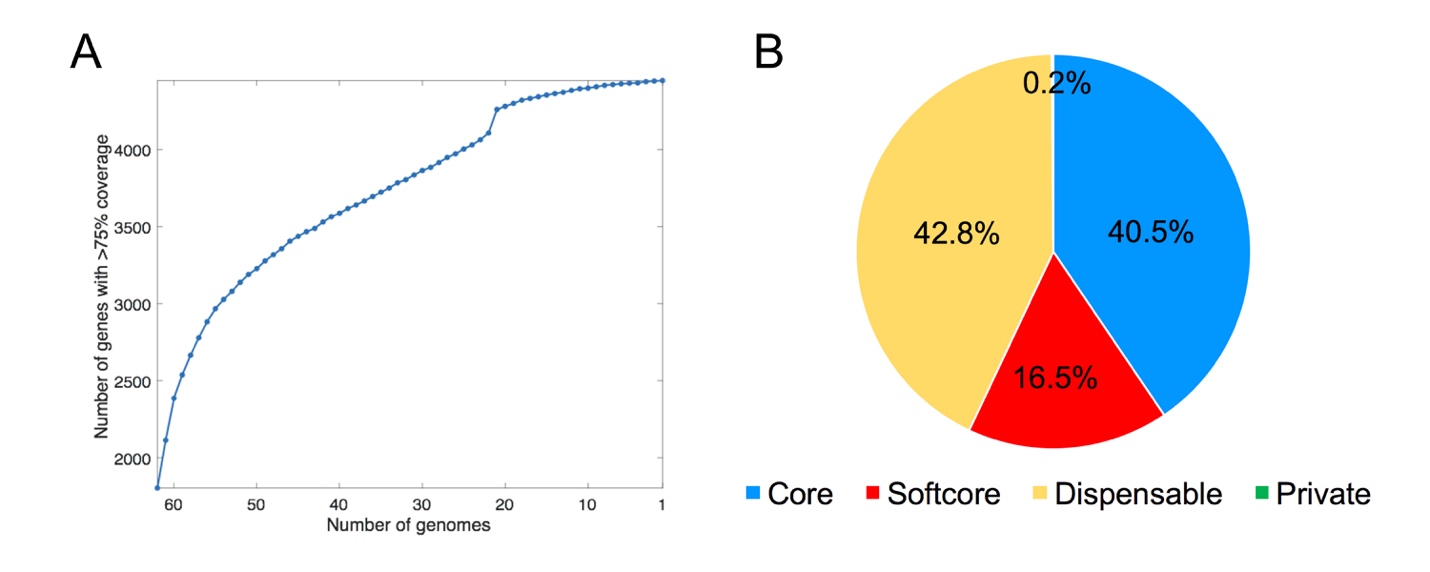

**Supplemental Fig. S6 Characterization of core non-coding genes.** (A) Cumulative curve of non-coding genes across 62 sorghum accessions. (B) Distribution of core, softcore, dispensable and private non-coding genes identified from 62 sorghum accessions.

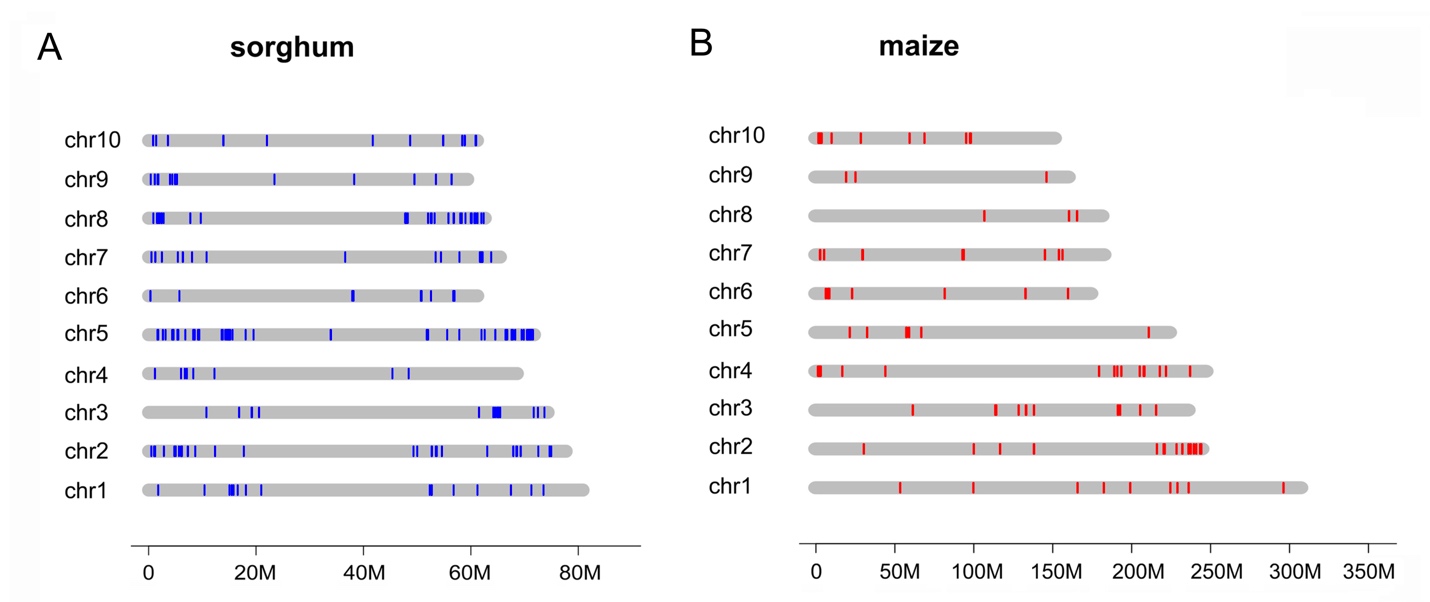

**Supplemental Fig. S7** Chromosome distribution of NBS-LRR genes in (**A**) sorghum and (**B**) maize.

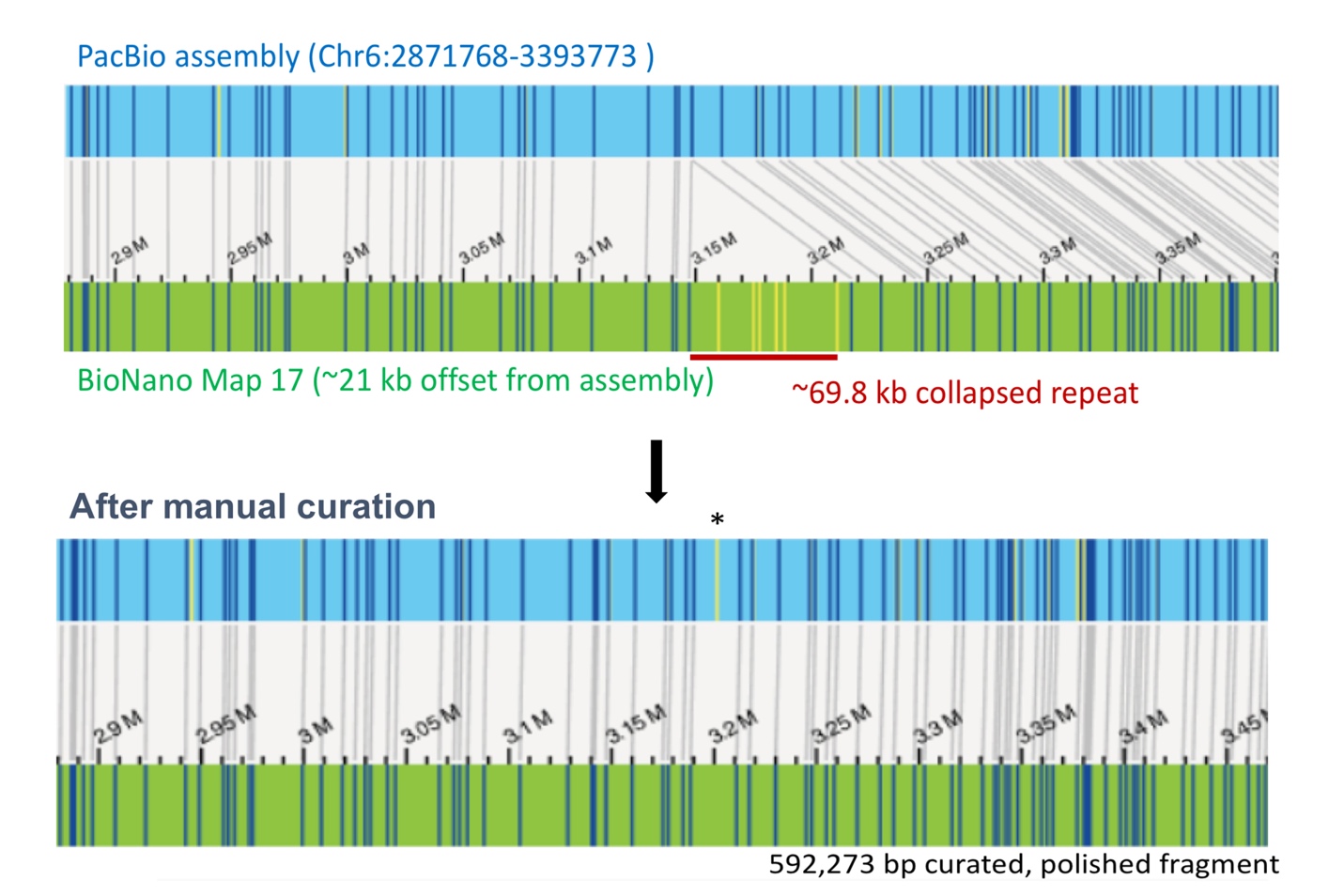

**Supplemental Fig. S8 Manual curation of assembly in the SCA region.**

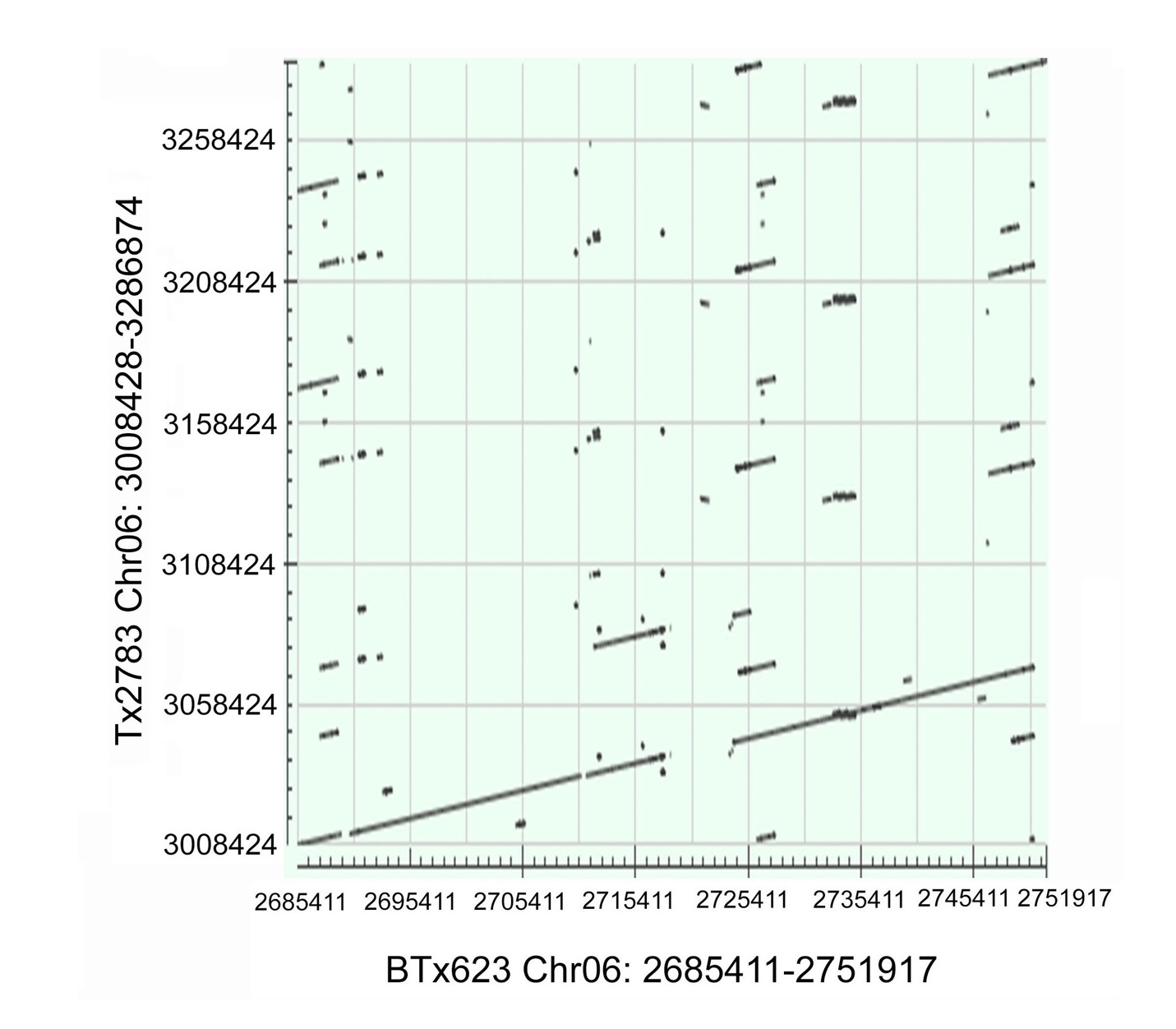

**Supplemental Fig. S9 Dotplot of deletions in the SCA resistance region between Tx2783 and BTx623.**

**Supplemental Tables**

**Supplemental Table S1. Summary of Bionano data for Tx2783, RTx436 and BTx623.**

|  | **Tx2783** | **RTx436** | **BTx623** |
| --- | --- | --- | --- |
| **Input molecule stats (filtered)** | | | |
| Total number of molecules | 198,889 | 432,088 | 361,453 |
| Total length (Mbp) | 139,053.311 | 196,107.731 | 176,721.480 |
| Minimum molecule size (kbp) | 550.000 | 350.000 | 350.000 |
| Molecule N50 (kbp) | 682.584 | 441.166 | 479.676 |
| Label density (/100kb) | 16.601 | 16.291 | 16.642 |
| **De novo assembly** | | | |
| Total Map Count | 27 | 28 | 30 |
| Maps per Chromosomes (nuclear genome) | 1.9 | 1.8 | 2.1 |
| Total Genome Map Length (Mbp) | 721.504 | 723.680 | 719.701 |
| Genome Map N50 (Mbp) | 36.987 | 37.781 | 36.133 |
| Largest Map (Mbp) | 86.486 | 87.280 | 86.174 |
| Total Sorghumv3.1 Reference Length (Mbp) | 732.152 | 732.152 | 732.152 |
| Total unique alignment to SorgumV3.1 (Mbp) | 396.709 | 388.604 | 680.886 |
| Effective coverage of assembly (X) | 148.228 | 141.249 | 163.812 |
| Average confidence | 37 | 29.6 | 51.2 |

**Supplemental Table S2. Summary of TE annotations for Tx2783, RTx436, and BTx623 genomes from EDTA.**

| **Class** |  | **Count** | | | **bpMasked** | | | **%masked** | | |
| --- | --- | --- | --- | --- | --- | --- | --- | --- | --- | --- |
| **DNA** |  | **Tx2783** | **RTx436** | **BTx623** | **Tx2783** | **RTx436** | **BTx623** | **Tx2783** | **RTx436** | **BTx623** |
|  | DTA | 12151 | 13551 | 10811 | 4002631 | 4300733 | 3714573 | 0.56% | 0.61% | 0.53% |
|  | DTC | 54956 | 78971 | 53061 | 25885444 | 27635943 | 26098836 | 3.65% | 3.92% | 3.74% |
|  | DTH | 75118 | 23076 | 75669 | 14838564 | 7065184 | 15329029 | 2.09% | 1.00% | 2.19% |
|  | DTM | 76210 | 37819 | 56303 | 12828965 | 9330520 | 10213875 | 1.81% | 1.32% | 1.46% |
|  | DTT | 76446 | 68909 | 61467 | 14137781 | 13739557 | 11825599 | 1.99% | 1.95% | 1.69% |
|  | Helitron | 44954 | 54592 | 54880 | 17111527 | 26593000 | 27922368 | 2.41% | 3.77% | 4.00% |
| **LTR** |  |  |  |  |  |  |  |  |  |  |
|  | Copia | 42287 | 54185 | 36194 | 47560408 | 45212812 | 42075707 | 6.71% | 6.41% | 6.02% |
|  | Gypsy | 322975 | 300150 | 284586 | 258933760 | 262519850 | 265302124 | 36.52% | 37.20% | 37.98% |
|  | Unknown | 136216 | 138739 | 145133 | 70903263 | 68284050 | 64212478 | 10.00% | 9.68% | 9.19% |
| **MITE** |  |  |  |  |  |  |  |  |  |  |
|  | DTA | 7010 | 6739 | 6203 | 1575047 | 1459219 | 1294434 | 0.22% | 0.21% | 0.19% |
|  | DTC | 1232 | 1167 | 1070 | 211606 | 200423 | 182619 | 0.03% | 0.03% | 0.03% |
|  | DTH | 36353 | 35689 | 39902 | 5480776 | 6309477 | 6193680 | 0.77% | 0.89% | 0.89% |
|  | DTM | 109602 | 110469 | 63445 | 13816736 | 14731404 | 8459241 | 1.95% | 2.09% | 1.21% |
|  | DTT | 10908 | 11914 | 10604 | 1679352 | 1894854 | 1625969 | 0.24% | 0.27% | 0.23% |
|  | Total interspersed | 1006418 | 935970 | 899328 | 488965860 | 489277026 | 484450532 | 68.97% | 69.34% | 69.35% |

**Supplemental Table S3. Status of protein-coding gene annotations for Tx2783 and RTx436.**

|  | Tx2783 | RTx436 | BTx623 community |
| --- | --- | --- | --- |
| gene count | 29612 | 29265 | 34118 |
| gene length(ave) | 3833 | 3900 | 3714 |
| gene length(median) | 2888 | 2917 | 2824 |
| exon count | 147196 | 146857 | 154042 |
| exon length(ave) | 357 | 353 | 449 |
| exon length(median) | 172 | 171 | 176.5 |
| intron count | 117584 | 117592 | 119924 |
| intron length(ave) | 512 | 514 | 454 |
| intron length(median) | 150 | 150 | 142 |
| CDS count | 35998 | 41713 | 47110 |
| CDS length(ave) | 980 | 838 | 842 |
| CDS length(median) | 1005 | 1002 | 981 |
| peptide count | 35998 | 41713 | 47110 |
| peptide length(ave) | 327 | 279 | 281 |
| peptide length(median) | 335 | 334 | 327 |
| five_prime_UTR count | 32455 | 30983 | 25100 |
| five_prime_UTR length(ave) | 197 | 196 | 484 |
| five_prime_UTR length(median) | 147 | 143 | 207 |
| three_prime_UTR count | 29444 | 29109 | 26660 |
| three_prime_UTR length(ave) | 371 | 374 | 653 |
| three_prime_UTR length(median) | 318 | 314 | 356 |
| exons per transcript (ave) | 5.0 | 5.0 | 4.5 |
| single-exon gene count (pct) | 6783 (22.9) | 6585 (22.5) | 8467 (24.8) |

*Note: longest_CDS_transcript_used_for_calculation_of_CDS_and_Protein_length_statistics

**Supplementary Table S4. The list of large inversions that detected in Tx2783 and RTx436.**

| Chromosome | Start | End | Length (bp) |
| --- | --- | --- | --- |
| Chr01 | 33356132 | 35275835 | 1,919,703 |
| Chr01 | 36401709 | 37214054 | 812,345 |
| Chr01 | 75514170 | 76287030 | 772,860 |
| Chr02 | 3525124 | 4442667 | 917,543 |
| Chr02 | 7129333 | 8080508 | 951,175 |
| Chr02 | 59278465 | 60164539 | 886,074 |
| Chr02 | 65098169 | 65956262 | 858,093 |
| Chr02 | 67337397 | 68529467 | 1,192,070 |
| Chr02 | 69546710 | 70398650 | 851,940 |
| Chr04 | 17591213 | 18931707 | 1,340,494 |
| Chr04 | 46159835 | 46971912 | 812,077 |
| Chr05 | 1040227 | 1305930 | 265,703 |
| Chr05 | 12537018 | 12913225 | 376,207 |
| Chr05 | 21415297 | 24383781 | 2,968,484 |
| Chr05 | 30915630 | 36344641 | 5,429,011 |
| Chr05 | 39957499 | 44105202 | 4,147,703 |
| Chr05 | 62357906 | 63977956 | 1,620,050 |
| Chr05 | 66202036 | 66402440 | 200,404 |
| Chr06 | 31936255 | 33531392 | 1,595,137 |
| Chr06 | 35972918 | 37471793 | 1,498,875 |
| Chr06 | 43255203 | 43523149 | 267,946 |
| Chr06 | 55583217 | 56160866 | 577,649 |
| Chr06 | 60667208 | 61180903 | 513,695 |
| Chr07 | 4044542 | 4935169 | 890,627 |
| Chr07 | 9100451 | 11053358 | 1,952,907 |
| Chr07 | 32761311 | 38838240 | 6,076,929 |
| Chr07 | 40289965 | 41528867 | 1,238,902 |
| Chr07 | 55533219 | 56081737 | 548,518 |
| Chr08 | 1217649 | 2011863 | 794,214 |
| Chr08 | 25886376 | 30628805 | 4,742,429 |
| Chr08 | 50775371 | 51218866 | 443,495 |
| Chr08 | 57617369 | 58494926 | 877,557 |
| Chr09 | 12233033 | 12383816 | 150,783 |
| Chr09 | 14584929 | 16080409 | 1,495,480 |
| Chr09 | 25853901 | 27504757 | 1,650,856 |
| Chr09 | 45499346 | 45780113 | 280,767 |

**Supplemental Table S5. Number of SNPs called in each genome relative to the Tx2783 reference.**

|  | **Tx2783** |
| --- | --- |
| **BTx623** | 2,851,709 |
| **RTx430** | 3,214,708 |
| **RTx436** | 2,991,813 |
| **Rio** | 2,897,208 |

**Supplemental Table S6. Number of SVs called in each genome relative to the Tx2783 reference.**

|  | **BTx623** | **RTx436** | **RTx430** | **Rio** |
| --- | --- | --- | --- | --- |
| Insertion | 37763 | 43142 | 47480 | 39154 |
| Deletion | 41293 | 42760 | 46897 | 44789 |
| CNV | 20592 | 22548 | 28469 | 30086 |
| INV | 4992 | 3288 | 4531 | 7047 |
| Total | 104640 | 111738 | 127377 | 121076 |

**Supplemental Table S7. Number of core, softcore, dispensable and private genes under various cutoffs of gene coverage for protein-coding and non-coding genes identified from 62 sorghum accessions.**

|  | 50% gene coverage cutoff  protein coding (non-coding) | 75% gene coverage cutoff  protein coding (non-coding) | 90% gene coverage cutoff  protein coding (non-coding) |
| --- | --- | --- | --- |
| core | 22110 (2498) | 17314(1803) | 11690(1140) |
| soft core | 3477(748) | 3466(734) | 3307(716) |
| dispensible | 5216(1210) | 9952(1906) | 15376(2550) |
| private | 102(3) | 173(7) | 532(44) |

**Supplemental Table S8. Number of NLR loci in each genome, identified by genome scanning.**

| **Genome** | **Number of NLR loci** |
| --- | --- |
| RTx430 | 363 |
| Tx2783 | 355 |
| RTx436 | 350 |
| BTx623 | 333 |
| Rio | 319 |

**Supplemental Table S9. Summary of SVs in the SCA resistance region.**

| **Tx2783**  **Chr** | **Tx2783**  **start** | **Tx2783**  **stop** | **SV** | **BTx623**  **Chr** | **BTx623**  **start** | **BTx623**  **stop** | **Size of SV (bp)** | **Overlapped gene in Tx2783** |
| --- | --- | --- | --- | --- | --- | --- | --- | --- |
| Chr06 | 3012626 | 3012643 | DEL | Chr06 | 2690273 | 2690274 | 17 | NA |
| Chr06 | 3013211 | 3013277 | DEL | Chr06 | 2690839 | 2690837 | 66 | NA |
| Chr06 | 3056117 | 3056126 | DEL | Chr06 | 2735297 | 2735298 | 9 | NA |
| Chr06 | 3057342 | 3057383 | DEL | Chr06 | 2736452 | 2736553 | 60 | NA |
| Chr06 | 3068938 | 3069191 | DEL | Chr06 | 2748046 | 2748296 | 253 | NA |
| Chr06 | 3080438 | 3080599 | DEL | Chr06 | 2713358 | 2713520 | 161 | NA |
| Chr06 | 3082064 | 3084412 | DEL | Chr06 | 2714983 | 2717317 | 2348 | SbiRTX2783.06G018300 |
| Chr06 | 3091201 | 3282740 | DEL | Chr06 | 2725441 | 2747807 | 191539 | SbiRTX2783.06G018400;  SbiRTX2783.06G018450; SbiRTX2783.06G018460; SbiRTX2783.06G018500; SbiRTX2783.06G018600; SbiRTX2783.06G018700. |
| Chr06 | 3283229 | 3284121 | DEL | Chr06 | 2748294 | 2749177 | 892 | SbiRTX2783.06G018800 |
| Chr06 | 3284749 | 3285059 | DEL | Chr06 | 2749803 | 2750109 | 310 | NA |
| Chr06 | 3285936 | 3286510 | DEL | Chr06 | 2750984 | 2751554 | 574 | NA |
